# Functional trade-offs: exploring the temporal response of field margin plant communities to climate change and agricultural practices

**DOI:** 10.1101/2023.03.03.530956

**Authors:** Isis Poinas, Christine N Meynard, Guillaume Fried

**Author notes:** Corresponding author: Isis Poinas. **Authorship statement:** I.P., G.F. and C.N.M. planned and designed the research; I.P. analyzed the data and wrote the first draft of the manuscript; G.F. and C.N.M. contributed substantially to revisions. All authors gave final approval for publication. **Data accessibility statement:** Data available via the Data INRAE Repository at https://doi.org/10.57745/QAMAWM.

## Abstract

Over the past decades, agricultural intensification and climate change have led to vegetation shifts. However, functional trade-offs linking traits responding to climate and farming practices are rarely analyzed, especially on large-scale empirical studies. Here we used a standardized yearly monitoring effort of agricultural field margin flora at the national scale to assess the temporal response of diversity and functional traits to variations in climate and intensity of agricultural practices. We examined temporal trends in climate (temperature, soil moisture), intensity of agricultural practices (herbicides, fertilization, margin management), plant species richness, and community-weighted means and variances of traits expected to vary both with climate and agricultural practices (e.g. seed mass, specific leaf area), across 555 sites in France between 2013 and 2021. We found clear temporal climatic trends (temperature increased while soil moisture decreased), whereas trends in agricultural practices were weak over the past decade, with only slight decreases in herbicides and margin management intensity. During the same period, functional changes in plant communities were significant, showing an increase of thermophilic species (including some Mediterranean species) with a conservative resource acquisition strategy (high stature, late and short flowering), mainly explained by climate change. The reduction in field margin management intensity (mainly mowing), also resulted in a vegetation shift towards a more conservative strategy. In contrast, there was no impact of the slight temporal changes of practices conducted within cultivated fields (herbicides, fertilization) on vegetation changes. Our findings suggest that species adapted to climate change (including Mediterranean and conservative species) have temporally increased in proportion. Importantly, we identified functional trade-offs indicating that these species are also the most vulnerable to intensive agricultural practices, as they are less adapted to high levels of resources and disturbances. We put these results into the conceptual framework of Grime’s CSR triangle and revealed a temporal decline of competitive and ruderal species in favor of stress-tolerant species better adapted to climate change. Choosing less intensive management can broaden the functional spectrum of agricultural plant communities, by maintaining the ability of stress-tolerant species selected by climate change to colonize habitats largely dominated by ruderals. Put together, these results suggest that climate change and agricultural intensification could have synergistic negative impacts on plant diversity in field margins, highlighting the need to study several biodiversity drivers at the same time in anthropized landscapes.

## Introduction

Since the 1950s, agricultural intensification has led to declining biodiversity (Emmerson et al., 2016), while climate change has caused notable changes in a wide range of taxa and habitats (Lovejoy, 2006). However, teasing out the relative importance of these two drivers on community trajectories can be quite challenging (Oliver & Morecroft, 2014). Since agricultural intensification took place in the 1950s in Europe, the main changes linked to agricultural practices (notably in terms of intensity in pesticide use and fertilization) in plant communities have likely already occurred (Lososová et al., 2004). For example, a meta-analysis considering 32 studies across Europe and covering the time period from 1939 to 2011 showed that weed species richness declined up to the 1980s, but has stabilized or even increased since then (Richner et al., 2015). Pesticide reduction plans have had so far little effect in France (Guichard et al., 2017), hindering the detection of temporal changes in biodiversity linked to changes in pesticide use. Conversely, short-term declines in species diversity due to climate change have been observed (e.g. Fonty et al., 2009), and recent temperature increases in France may impact plant communities similarly (Baude et al., 2022; Martin et al., 2019).

Temporal changes in plant communities cannot be discerned solely using taxonomic approaches due to the differing traits affected by resource availability and disturbance levels (Garnier & Navas, 2012); therefore, a functional approach provides an additional perspective to accurately understand these changes. This is particularly important in agroecosystems, where both resource (fertilization) and disturbance (herbicides, field margin management) gradients play crucial roles in shaping communities (Gaba et al., 2014; MacLaren et al., 2020). For instance, weeds with a ruderal strategy (low height and seed mass, long and early flowering, high SLA) are better adapted to agricultural disturbances, such as tillage, herbicides or management by mowing (Grime, 2006; Fried et al., 2022). At the same time, traits responding to agricultural practices can co-vary with other traits that are linked to resource acquisition, competitive ability, or climate. For example, seed mass, which is often used as a proxy for competitive ability, increases along soil fertility, temperature and solar radiation gradients (Fried et al., 2022; Murray et al., 2004). Furthermore, correlations among different traits may represent trade-offs that impact community adaptation (Díaz et al., 2016; Wright et al., 2004). Because of these trade-offs, selective pressures due to one driver (e.g. climate change) may impact community properties along the other (e.g. agricultural intensification). In this context, Grime (1977) proposed a framework called the CSR triangle, which defines three strategies (competitiveness, stress-tolerance, and ruderality) along two axes of variation (resource and disturbance). These strategies are correlated to multiple traits and have proven useful to understand plant community dynamics in agroecosystems (Fried et al., 2022). As traits responding to climate and agricultural practices may co-vary (Garnier & Navas, 2012), it can be difficult to identify the main drivers behind community changes, making this framework potentially useful to understand such trade-offs.

To understand the complex interactions between climate change and agricultural practices, it is thus essential to examine temporal changes in species trait distribution. For example, in French wheat fields, ruderal species increased their frequency between the 1970s and 2000s, potentially due to their ability to escape recurrent disturbances, such as herbicide applications (Fried et al., 2012). Inter-annual variations in specific leaf area, leaf dry matter content and plant height are related to nitrogen supply (Borgy et al., 2017; Gaba et al., 2014), while increased precipitations push the foliar economic spectrum towards more acquisitive species (i.e. with higher SLA; Wheeler et al., 2023). Additionally, mean thermal preference of plant communities, as well as their phenology, can vary over time in response to temperature changes, even over relatively short periods (Bellard et al., 2012; Martin et al., 2019). These temporal variations in functional traits reveal patterns that cannot be assessed solely with space-for-time substitution.

In this study, we aimed at deciphering how inter-annual temporal variations and temporal trends in climate (temperature, soil moisture) and agricultural practices (frequency of herbicide use, margin management and nitrogen dose in fertilizers) in France structure species richness, trait composition and ecological strategies of field margin plant communities. We studied the herbaceous field margin, which represents the uncultivated vegetated area located between the cultivated strip and the adjacent habitat. Using a standardized national monitoring effort spanning 9 years (2013-2021) in 555 agricultural field margins covering continental France, our study stands as one of the first to investigate the temporal trends in agricultural practices and climate, and explore the response of species richness, trait composition and ecological strategies to these trends at such extensive scales. We hypothesized that plant traits sensitive to temperature and soil moisture will co-vary with temporal warming trends while agricultural practices would have a comparatively weaker temporal influence on plant communities, as we did not expect clear temporal trends in these practices. We also expected a limited impact of agricultural practices on margin plant communities, because field margins only receive a small amount of nitrogen and herbicides drifting from neighboring plots. Furthermore, we explored the connection between Grime’s CSR strategies, climate and farming practices. Considering that these strategies are linked to resource and disturbance levels, we hypothesized that they would respond to climate factors (particularly reduced soil moisture) and agricultural practices (disturbance and resource provision through fertilization). On top of the national analyses, and because this dataset includes the Mediterranean flora, which has been shown to respond more strongly to some agricultural filters (Poinas et al., 2023), we included analyses separating this region from the rest of France. We also separated vineyards from annual crops, because vineyards include very different management practices and no crop rotation (Metay et al., 2022). Finally, we also analyzed annual plant species separately, as they may respond more rapidly to environmental changes (Martin et al., 2019; Fitter & Fitter, 2002).

## Materials and methods

### Vegetation surveys

We used vegetation data from the 500-ENI network, which is funded by the French Ministry of Agriculture (see details in Andrade et al., 2021) and monitored 555 agricultural field margins across continental France between 2013 and 2021 (with some site turnover) (**Fig. 1**). These survey sites represented three main crop types **(Appendix A, Fig. SA. 1)**: annual crops (with winter wheat or maize as the main crop production in the rotation), market gardening crops (mainly lettuce) and vineyards. The proportion of sites under organic farming was roughly 20%, but agricultural practices covered a wide range of pesticide application, fertilizers and soil management. Within each survey site, plant species were identified in ten 1 m² quadrats along the field margin (**Appendix A**, **Fig. SA.2**). Presence-absence of each species was recorded for each quadrat, which provided a frequency of occurrence from 0 to 10 in each field margin, used here as an index of relative abundance. Surveys were performed once per year at peak flowering (between the end of April and the beginning of August, depending on the region). At the national scale, this represented 4172 observations (year x site), leading to the identification of 852 taxa. Because observers changed among sites and over time (312 observers in total, each observer following on average 5 distinct sites during 4 years) and did not have the same level of expertise, we constrained our analyses to a subset of 142 focal species (Andrade et al., 2021) which are expected to be known by all the observers (and thus removing 11% of the total abundances).

**Fig. 1.**
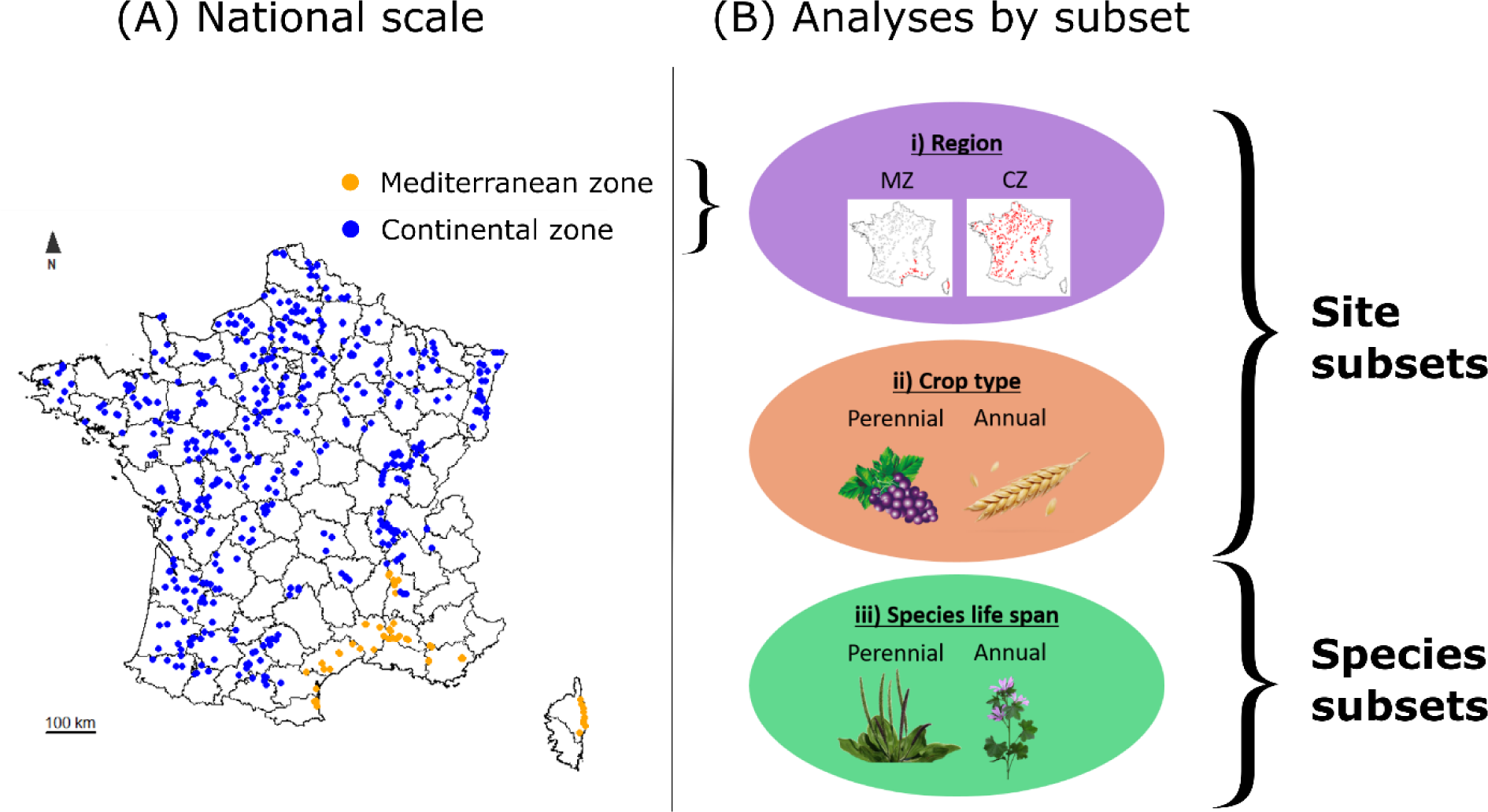
(A) Distribution map of the 555 field margins monitored at least one year between 2013 and 2021 in France. The black lines represent the limits of French departments. Orange: sites in Mediterranean zone (n = 57), blue: sites in Continental zone (n = 498). The contours of the Mediterranean zone (MZ) were derived from the Mediterranean zone and Corsica as defined in the VégétalLocal map (Office français de la biodiversité, 2021); the rest of France will be referred to here as Continental zone (CZ). (B) Subsets of data used in additional analyses: i) the regional scale splits the MZ from the CZ; ii) annual crops included rotations based on wheat, maize and market gardening crops (n = 450); perennial crops only included vineyards (n = 105); iii) annual plants (n = 61) opposed to perennials (n = 79).

### Climatic and agricultural variables

We gathered two types of explanatory variables: the first came directly from the 500-ENI network and reflects agricultural practices assessed directly on the monitoring sites; the second one included meteorological data from an external database (see below). Here, we chose not to include landscape factors, as a previous study on the same dataset demonstrated that landscape variables account for a negligible proportion of variance at the national scale, in contrast to climate (Poinas et al., 2023).

Agricultural practices were reported yearly from interviews of farmers into a standardized online database. Data collected relate to fertilization, herbicide use and field margin management (mainly mowing of vegetation). Daily meteorological data were extracted from the SAFRAN climate model of Météo France, with a resolution of 8 km (Le Moigne, 2002). Meteorological data were averaged over a one-year window prior to each floristic observation, while agricultural data were summed over the same period (**Table 1**). We selected variables that were weakly correlated (Spearman correlation < 0.65, **Appendix B**) and have been shown to influence plant communities in previous studies (**Table 1**, see **Appendix C** for the choice of variables).

**Table 1.**
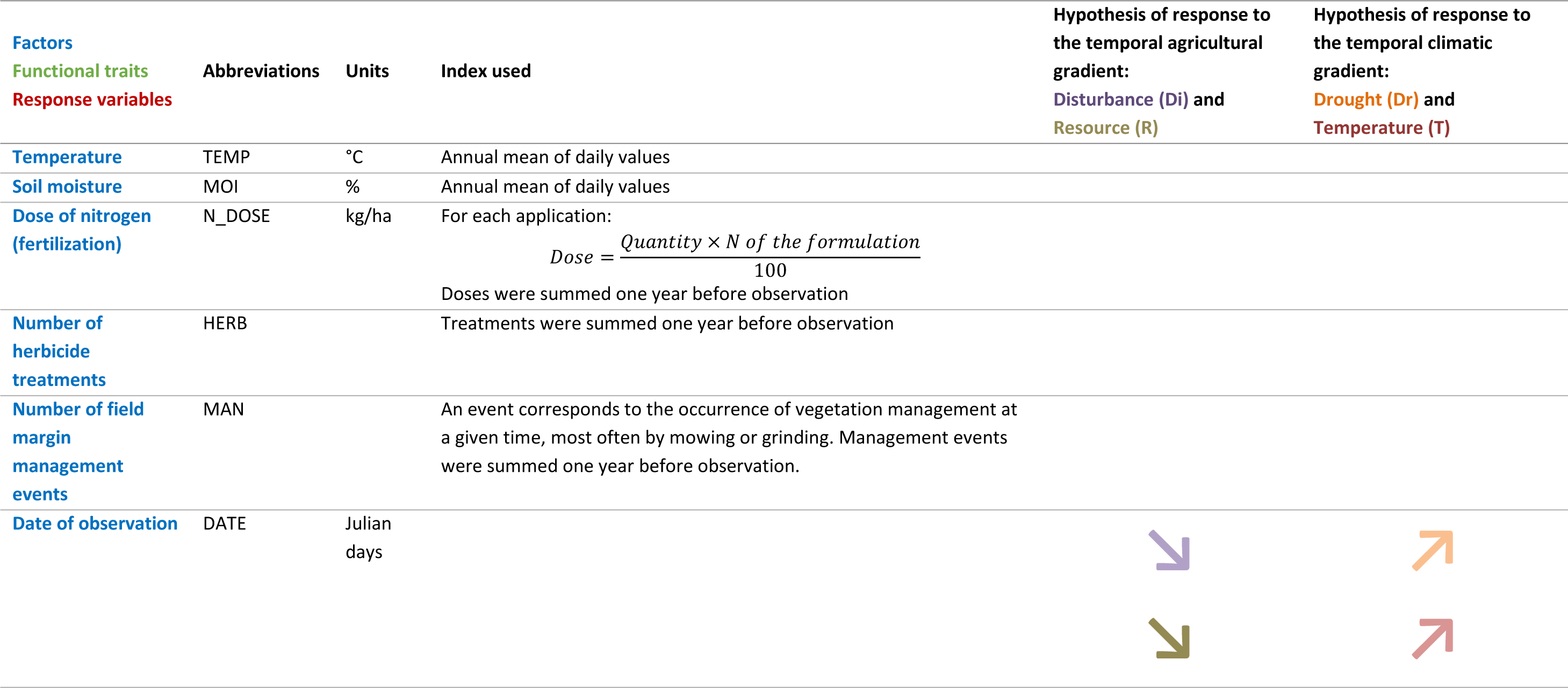
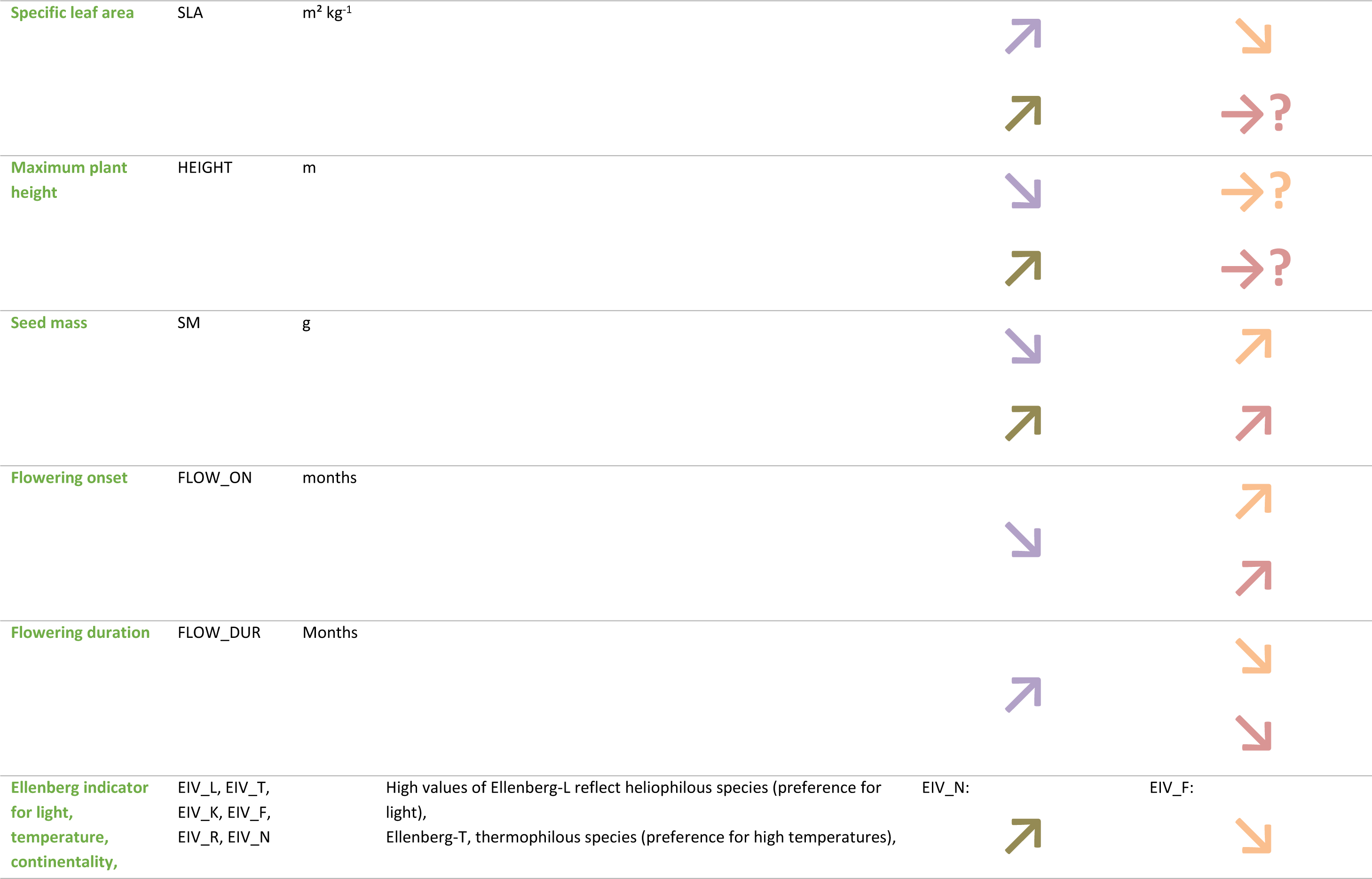
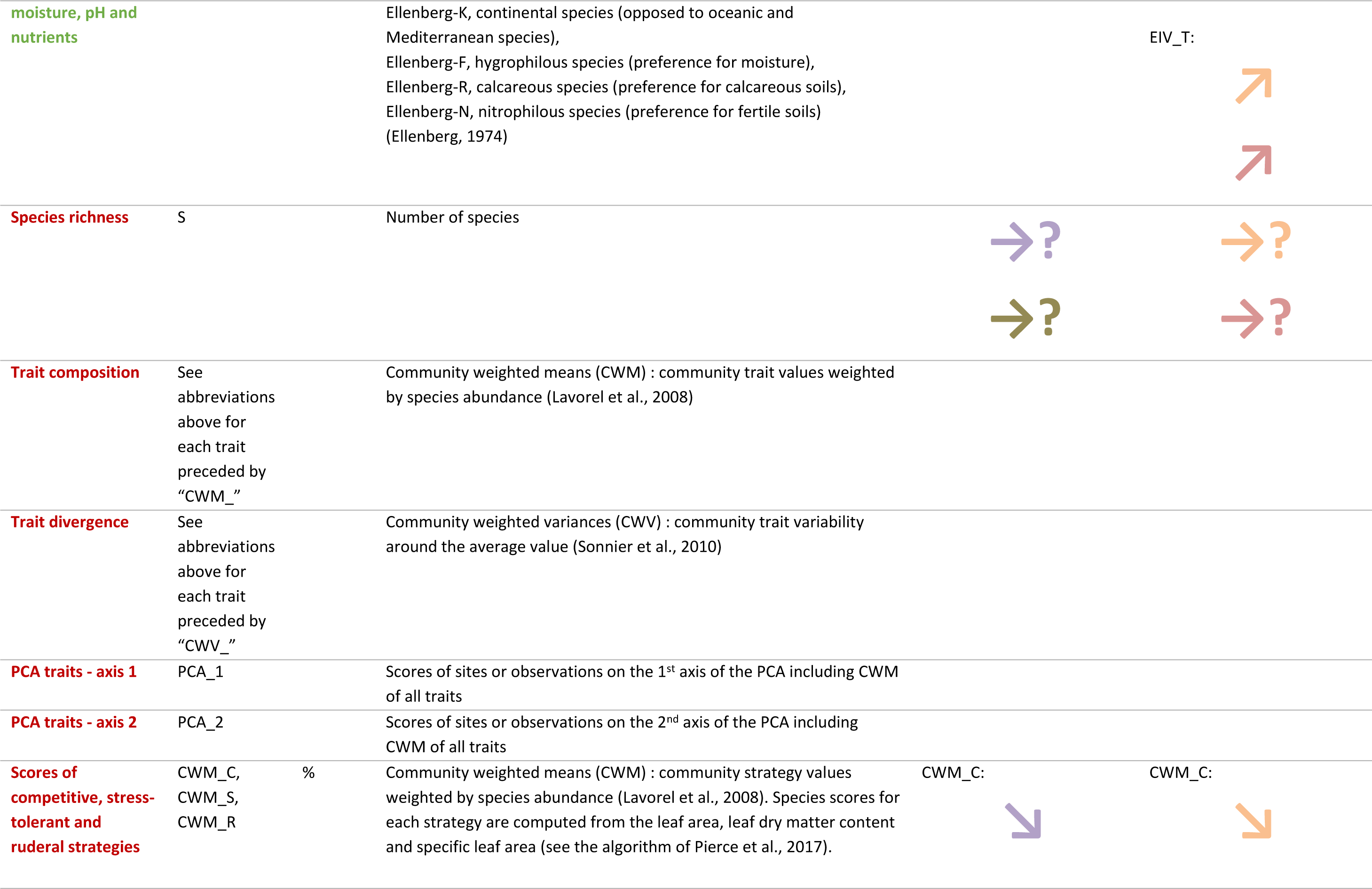
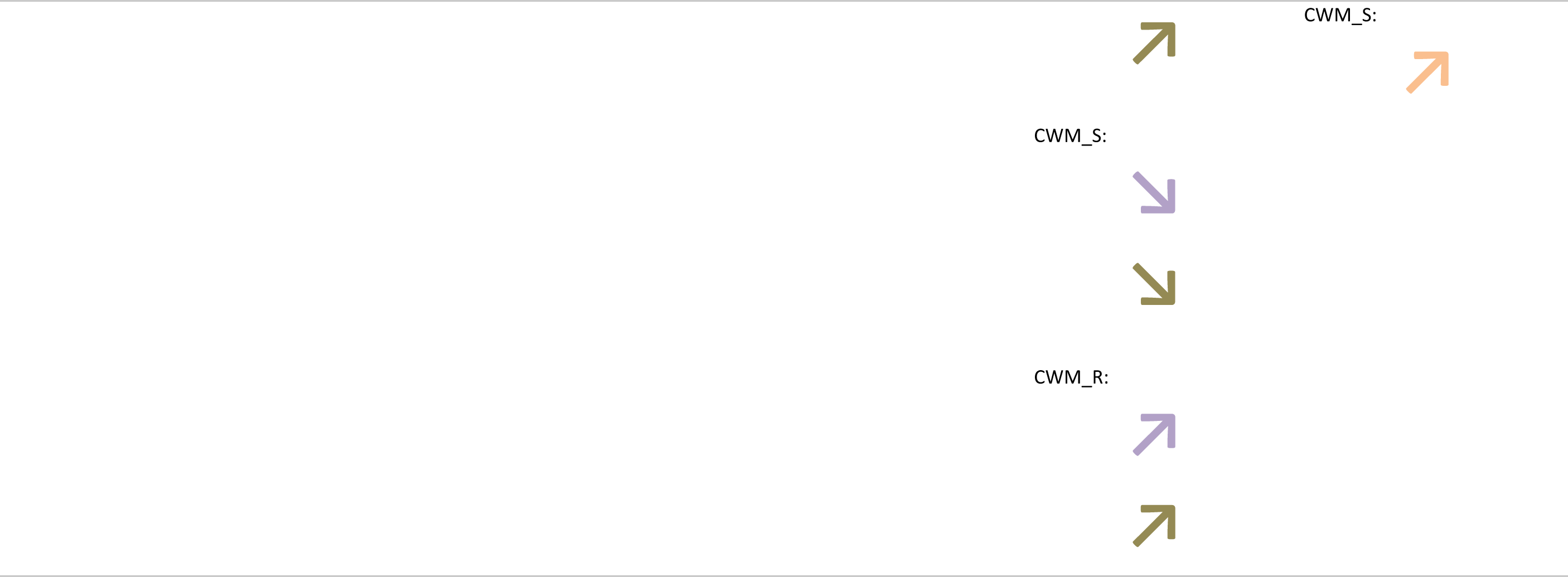
List of explanatory factors (blue), functional traits and ecological requirements (green) and response variables (red) with their abbreviations, units and calculation. We have illustrated by arrows the expected link of each factor and trait to the agricultural resource (fertilization) and disturbance gradient (herbicides and margin management), and to the climatic gradient (drought and increasing temperature). We also used arrows to illustrate the expected direction of variation of these gradients within a year (i.e. climatic and agricultural changes according to the date of observation). Horizontal arrows indicate contradictory findings in the literature (see **Appendix C** for the references).

### Plant functional traits

We extracted from external databases five functional traits and six species-level indices of ecological requirements (i.e. Ellenberg values; Ellenberg, 1974), assumed to respond to agricultural or climatic factors (**Table 1**, **Appendix B-C**). Functional traits were missing for four species, two of which could be imputed from an average over other species of the same genus. The remaining two species were removed from the analysis (representing 0.01% of the total abundances among the 142 species considered).

To characterize plant communities, we calculated species richness, community-weighted means (CWM) and community-weighted variances (CWV) of traits and ecological requirements for observations with at least three species (59 out of 4172 observations were excluded). The computation was performed using the R v.4.0.0 package _FD_, function *dbFD* for CWM, with the following formulas:

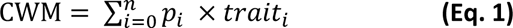

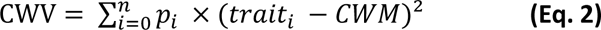

Where *p*_*i*_ is the relative abundance, *trait*_*i*_ is the value of trait for species *i,* and *n* is the total number of species. To correct for correlation between CWV and species richness, we used a null model approach, shuffling the abundances in the species matrix for species of the species pool, while keeping the species x trait matrix unchanged (Bopp et al., 2022). The species pool was defined by the site, allowing us to investigate temporal variations. This procedure keeps trait correlations, species richness and total abundance in a site unchanged, while dissociating abundances from trait values (Bernard-Verdier et al., 2012). To quantify the difference between observed and null CWV, we computed effect sizes (**Appendix D**). A positive effect size denotes a divergence in trait values within the community (convergence for negative effect size). These effect sizes (and not the raw CWV) were used in our analyses and referred to as CWV in the subsequent sections. We performed a normed PCA on the CWM of traits to classify each community based on its average trait combination or ecological strategy, which is reflected by its position on the first two axes.

### Plant functional strategies

According to Grime (1988), stress (i.e. a shortage of resources such as nutrients, water and light) and disturbance (i.e. the partial or total destruction of plant biomass) determine three main plant strategies representing combinations of traits that are viable under conditions of low disturbances and high resources (competitor, C), low disturbances and low resources (stress-tolerant, S) or high disturbances and high resources (ruderal, R). Originally developed to classify individual plant species into strategies, Grime’s theory can be useful to interpret functional changes in plant communities, especially in the context of global changes where vegetation is subject to harsher climatic conditions (more droughts) and various levels of agricultural disturbances.

To assess these strategies, we extracted the CSR scores for 119 out of 142 focal species from Pierce et al. (2017). CWM of CSR scores were computed by observation and were added to the PCA on the CWM of traits as supplementary variables. They were plotted on a CSR triangle to illustrate temporal trends in strategies.

### Temporal analyses of plant communities

The general framework of analyses is presented in **Fig. 2**. First, we checked if there was a temporal trend on the raw variables, and then we used climate and agricultural practices as predictors for the different response variables. In all cases, we used generalized additive mixed models (GAMM) to account for repeated measures at a site, with a Gaussian distribution in most cases (but see **Appendix E**, **Table SE.2**), and site identity as a random effect. Observer bias was accounted for by including the observer identity as a random term nested within sites. For each response variable (species richness, CWM, CWV and CSR strategies) and explanatory factor (temperature, soil moisture, nitrogen dose, herbicides and margin management), we built a first model with the year as a linear fixed effect. Then, a second model was built for each response variable, where climate, agricultural practices and observation date were linear explanatory factors. A first-order temporal autocorrelation structure within sites was included (Box et al., 2015). We removed observations with missing values in climatic and agricultural factors (1805 out of 4172 observations), and a few observations that distorted trait distributions (**Appendix E**), resulting in varying observation numbers across models (see **Fig. 5**). We repeated this analysis on subsets of data, including Mediterranean (MZ) vs Continental zones (CZ), margins adjacent to annual crops vs vineyards, and annual vs perennial plant species (**Fig. 1**). For all analyses, we chose a p-value threshold of 0.01 to focus on the effects for which our confidence level was highest.

**Fig. 2.**
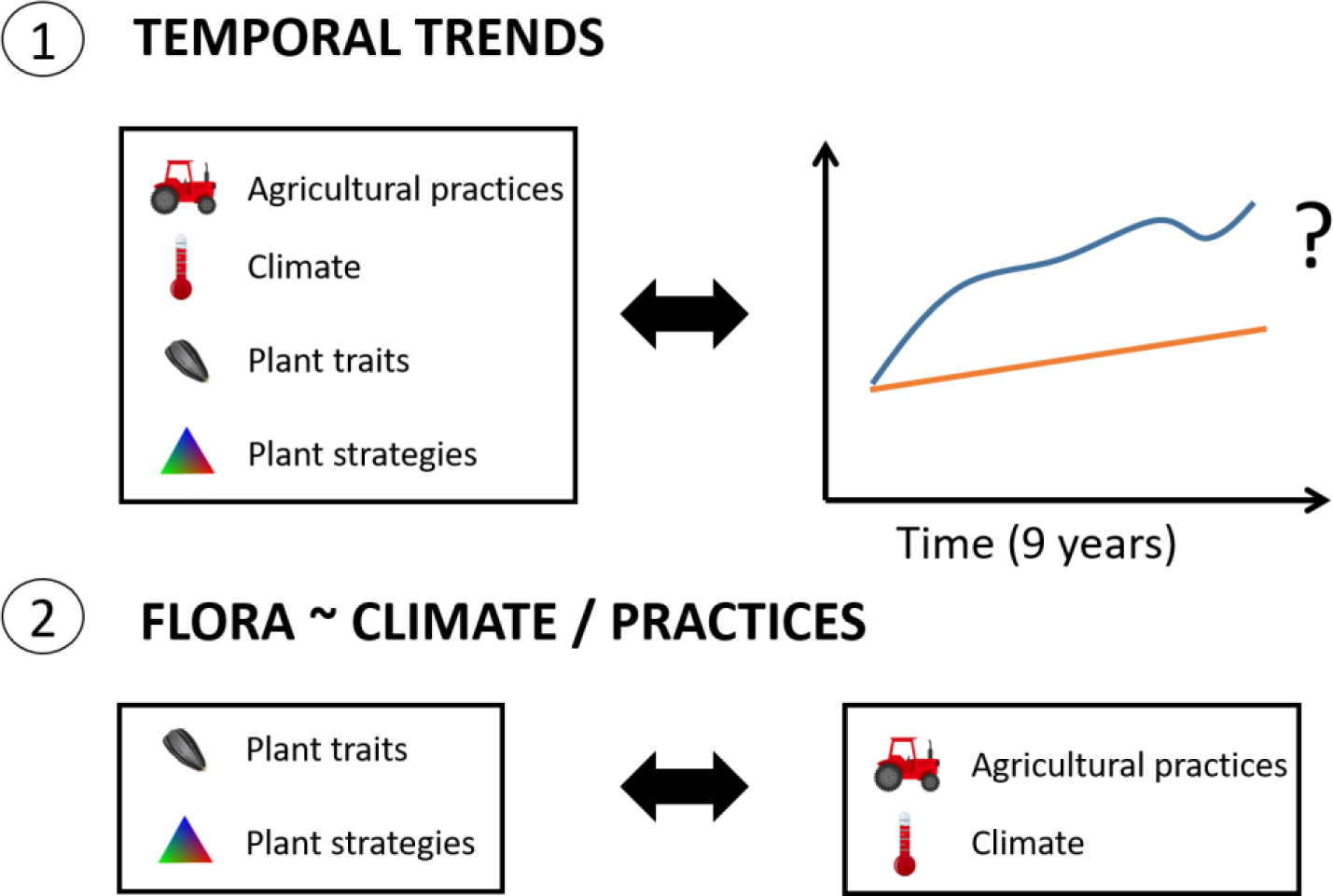
General framework of our analyses involved two main steps. (1) Firstly, we investigated the presence of any temporal trends in species richness, trait composition, ecological strategies, climatic and agricultural factors, as this provides crucial insights for the subsequent analysis. Indeed, given that we expected minimal temporal trends in agricultural practices, we did not expect significant temporal changes in flora in response to practices. (2) Secondly, we explored how plant communities have responded to the temporal changes in climate and agricultural practices.

## Results

### Temporal trends in climate, agricultural practices and plant communities

Temperatures have significantly increased by an average of 1.2°C over a decade (0.7°C in the Mediterranean Zone), while soil moisture has steadily declined (−14.1% by decade) (**Fig. 3**, and **Appendix F**). These trends differed between the Mediterranean zone (MZ) and the Continental zone (CZ), with the MZ experiencing a slower decline in soil moisture due to a high cumulative precipitation in 2019 (**Fig. 3**). Regarding agricultural practices, herbicides slightly decreased over time in vineyards (−0.9 application by decade; **Fig. 3**), with an even weaker trend in annual crops (−0.2 application by decade). Fertilization showed no significant temporal trend, except in vineyards where the cumulative dose of nitrogen has recently slightly increased (**Fig. 3**). The number of margin management events has decreased and particularly in the MZ (−0.5 by decade). Floristic surveys were conducted increasingly earlier in the season in the CZ (10.4 days earlier by decade) (**Appendix F**). Overall, there was a clear warming and drying trend in climate, but agricultural trends were weaker.

**Fig. 3.**
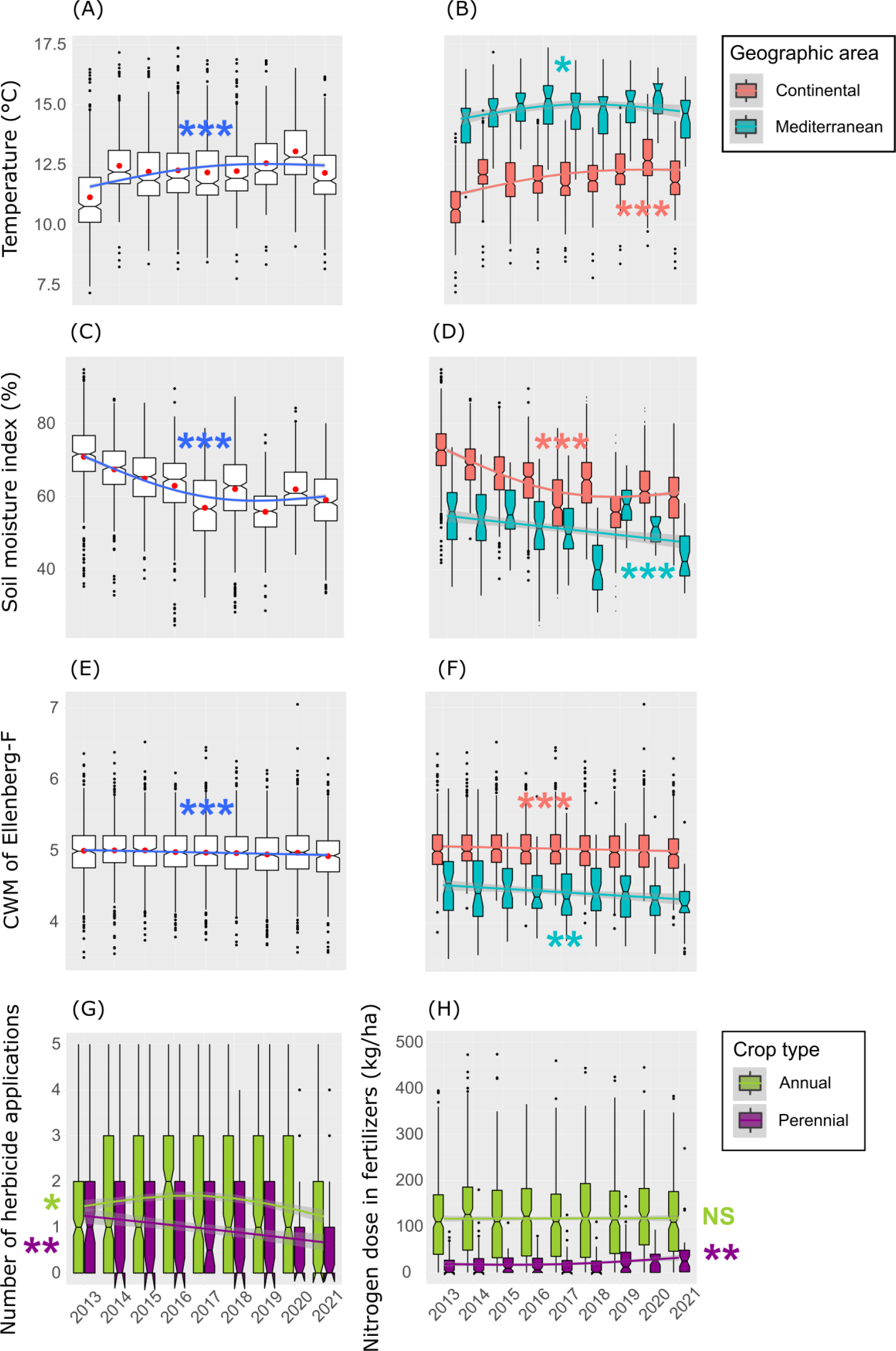
Temporal changes in temperature, soil moisture, CWM of Ellenberg-F (moisture requirement), number of herbicide applications and nitrogen dose in fertilizers. Red dots represent mean values. The curves are from a GAM, with a smooth term on the year restricted to three effective degrees of freedom. (A, C, E) National trend. (B, D, F) Trend by geographic area: CZ and MZ. (G, H) Trend by crop type: annual (wheat, maize, lettuce) and perennial (vineyard). Significance of smooth terms is referred as following: NS p ≥ 0.05; * p < 0.05; ** p < 0.01; *** p < 0.001.

Plant species richness has slightly increased over time at the national scale (+0.1 species by decade, **Fig. 5**), even more in the MZ (+0.4 species by decade) and vineyards (+0.3 species by decade) and only for annual species (**Appendix H**). We saw an increase in the CWM of maximum height (+5.8 cm by decade), seed mass (+0.2 g by decade), flowering onset (+3.1 days by decade) and a decrease in flowering duration (−7.8 days by decade, **Fig. 5**). The requirements for light, temperature and pH have increased, while those for moisture and nitrogen have declined. CWV (i.e. computed by comparison with expected CWV in a community of same richness) have decreased for most traits indicating trait convergence, and particularly for phenological traits such as flowering onset and flowering duration (−3.6 and −2.6 days by decade respectively), while they have increased for the requirements for temperature, pH and continentality, indicating trait divergence.

Changes in functional traits were more pronounced in the MZ, particularly for the flowering onset (+8.8 days by decade) and duration (−18.9 days by decade; **Appendix F**). Conversely, changes in Ellenberg values (environmental requirements) were only significant in the CZ and in annual crops. One exception was the temperature (Ellenberg-T) and moisture (Ellenberg-F) requirements, which have significantly changed in both the MZ and CZ. Interestingly, functional traits (and not environmental requirements) showed a temporal trend mainly for annual species (**Appendix F**).

### Functional trade-offs

The first PCA axis (named thereafter stress-tolerance axis, see **Appendix G, Fig. SG.1** for the correlation of each CSR strategy with each axis) explained 29.9% of the variation in traits and revealed a gradient from continental hygrophilous communities (high Ellenberg-K and F) associated with moist and resource-rich environments (high Ellenberg-N), to Mediterranean xero-thermophilous stress-tolerant communities (high Ellenberg-T and L, low Ellenberg-F) adapted to warm, arid and resource-poor environments (**Fig. 4, Appendix G, Fig. SG.2**). Communities with continental species were more nitrophilous (high Ellenberg-N), while Mediterranean communities had a higher seed mass. The second PCA axis (named thereafter ruderal axis) explained 19.5% of the variation and contrasted stress-tolerant/conservative communities adapted to low disturbance (low SLA, high stature, late and short flowering) with ruderal/acquisitive communities adapted to high disturbance (high SLA, short stature, early and long flowering).

**Fig. 4.**
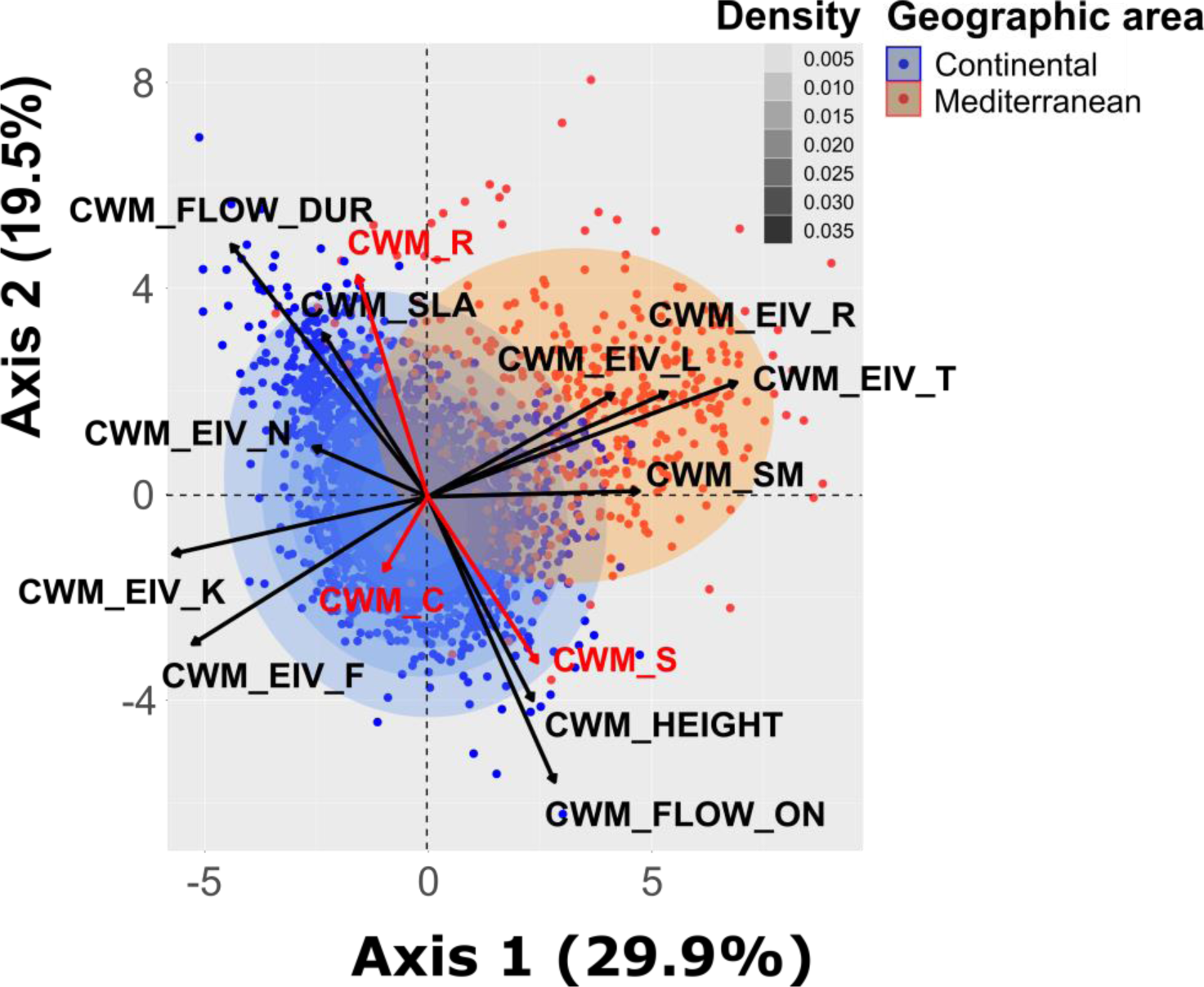
Normed PCA on CWM (first two axes) of functional traits and ecological requirements computed by observation. The color of the dots indicates the region to which they belong and the density curve highlights the concentration of data points in a given area. The correlations of traits to the PCA axes are in **Appendix G**, **Fig. SG.1** and the PCA for annual and perennial species in **Fig. SG. 3**. The CWM of strategies (in red) were added as supplementary variables. Abbreviations for CWM: CWM_SLA, specific leaf area; CWM_HEIGHT, maximum plant height; CWM_SM, seed mass; CWM_FLOW_ON, flowering onset; CWM_FLOW_DUR, flowering duration; CWM_EIV_L/T/K/F/R/N, requirement for light/temperature/continentality/moisture/pH/nitrogen; CWM_C, competitive strategy; CWM_S, stress-tolerant strategy; CWM_R, ruderal strategy.

### Temporal analyses of plant communities

Climatic factors were the predominant drivers of changes in community trait composition, with high R² for the temperature requirement (R² = 0.33) and stress-tolerance axis (see previous section, R² = 0.27, **Fig. 5**). Associations between each Ellenberg value and climatic factors opposed in a consistent way Mediterranean communities to nitrophilous continental ones along the stress-tolerance axis (**Fig. 5**). Increasing temperature increased the CWM of seed mass and Ellenberg-T (requirement for temperature) and decreased the CWM of SLA. Increasing temperature led to more divergence in the requirement for temperature and moisture (compared to a community of same richness). Conversely, increasing soil moisture brought convergence in the requirement for temperature, continentality and soil pH. Increasing temperature and drought were also associated with shorter flowering duration (−2.2 days by °C and +0.15 days by % of soil moisture), and later flowering onset only for increasing temperature (+1.6 days by °C; **Fig. 5**).

**Fig. 5.**
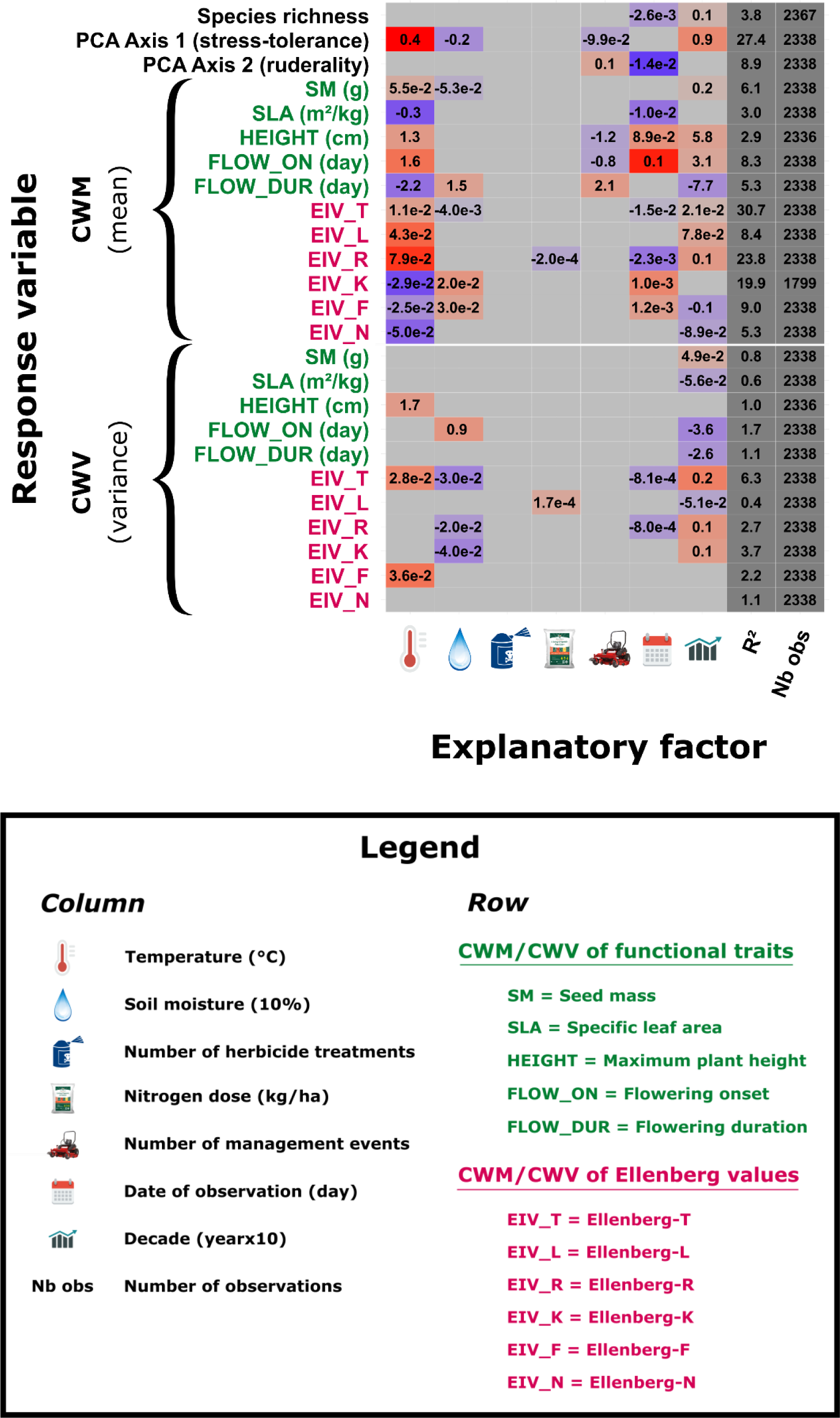
Results of temporal models (GAMM) on the whole dataset, with response variables in rows and explanatory factors in columns. The adjusted R², expressed as a percentage of variation, and the number of observations are reported. Significance is indicated by colored cells, with a p-value threshold of 0.01. Positive estimates are in red, negative estimates in blue, and the strength of the relationship (based on the standardized estimates) is reflected by the lightness of the color (weaker when lighter). It is important to note that the strength of the relationship can only be compared among explanatory factors for a same response variable. Reported values are the raw estimates and can be interpreted in the units of response and explanatory variables (e.g. an increase of 1°C in temperature leads to an increase in 1.6 days in the CWM of flowering onset). Models with the year as explanatory factor were run separately.

Margin management was the agricultural practice with the largest impact on changes in community trait composition, with an increase in its frequency associated with more ruderality (−1.2 cm in maximum height, -0.8 days in flowering onset and +2.1 days in flowering duration by management event). The date of observation also influenced changes in community trait composition, as a later observation was related to more conservative, competitive and continental communities, and to a decrease in species richness. Changes in the frequency of herbicide treatments had no significant effect, while an increasing annual nitrogen dose in fertilizers only slightly decreased the pH requirement (**Fig. 5**).

Results differed depending on the subset of data used (**Fig. 1**). In vineyards and the MZ, changes in soil moisture did not have any influence on changes in species richness or community trait composition (**Appendix H**) and temperature only increased the requirement for temperature (Ellenberg-T) and decreased SLA. When margins were more frequently managed in the MZ, Mediterranean species declined (decrease of temperature requirement and convergence towards higher values of continentality, **Appendix H**). Increasing nitrogen dose tended to decrease the number of species in the MZ, an effect also found on annual species. In vineyards, no agricultural effect was detected at the national scale. Annuals were more impacted by climatic variations and seasonal effects (observation date) than perennials, with high R² for temperature (R² = 0.39) and moisture (R² = 0.32) requirements.

## Discussion

Our study is one of the first to provide empirical evidence that climate change is already resulting in detectable functional changes in plant communities over a relatively short time interval of 10 years (see also Martin et al., 2019) (**Fig. 6**). Climate change tended to favor the stress-tolerance strategy at the expense of ruderality. These contrasting strategies highlight the functional trade-offs that prevent field margin plants from simultaneously adapting to climate change and intensive agricultural practices. Interestingly, reducing the frequency of margin management mimicked the impact of climate change on community trait composition, although the trend was less pronounced. Practices applied in the adjacent agricultural fields, including herbicide use and fertilization, had almost no effect on changes in community trait composition.

**Fig. 6.**
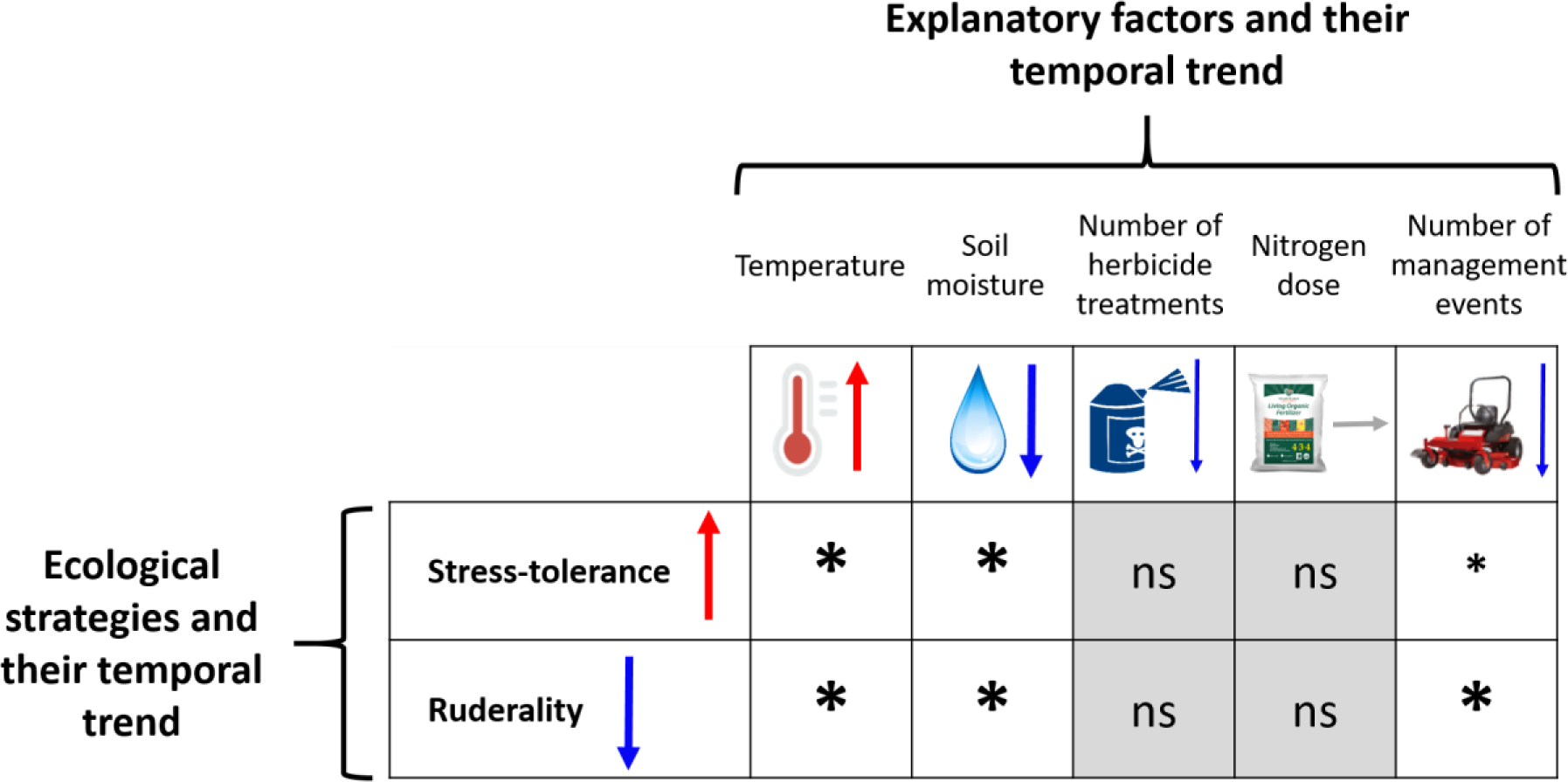
Synthesis of results based on temporal trends and temporal analyses of community response to changes in climate and practices. Ecological strategies are based on PCA axes (Axis 1 for stress-tolerance and Axis 2 for ruderality) and their associated traits. Explanatory factors and ecological strategies are depicted with their temporal trend over a decade (arrows). The asterisks illustrate the links between temporal trends in climate and practices and the resulting trends in communities, inferred from the coefficients in **Fig. 5** (ns = non-significant). The size of the asterisks represents the strength of the relationship, estimated from the number of impacted traits and standardized estimates in models.

### Climate as the main driver of temporal variations in field margin plant communities

Our analyses revealed a temporal shift towards more stress-tolerant and less ruderal communities, primarily driven by climate (Díaz et al., 2016; Pakeman et al., 2009). Since meteorological variables were extracted at a 8 km resolution, changes in soil moisture can be confidently attributed to climate change and not to the effect of soil compaction due to cultivation. Increasing temperature and drought favored more xerophilous and thermophilous species, with higher seed mass and lower nitrogen requirement, thus shifting the position of communities along the stress-tolerance axis. Our results also indicated that sites increasingly warmer and drier allowed for coexistence of a wider functional set of species, suggesting an increased abundance of thermophilous species without any loss of cold-adapted species. The increase in species richness over time provided additional support for this hypothesis.

The increase in temperature requirement at the community-level with rising temperatures was already documented, but mainly by studies covering entire floras (regional or local species pool) and time scales of several decades to a century (Salinitro et al., 2019; Tamis et al., 2005). We found that this trend is now detectable over a short-term period of only nine years (Martin et al., 2019). Interestingly, as in other recent studies (Duchenne et al., 2021; Martin et al., 2019), this trend was more pronounced in northern France, while Mediterranean communities responded less to climate change (**Appendix H**). On top of the fact that climatic trends observed in the MZ were weaker than in the CZ, Mediterranean species are already adapted to drought and heat stress, and might be more resilient to extinction risks (Thuiller et al., 2005). Because of their geographic position north of the Mediterranean Sea, they might also experience some competitive release due to the lack of immigrants coming from the south, and the northward shift of more temperate species (Duchenne et al. 2021).

Beyond the increase in temperature requirement, our models revealed additional temporal changes related to climate change that align well with the existing literature, including a decrease in mean SLA and an increase in mean seed mass and maximum height (Alarcón Víllora et al., 2019; Kühn et al., 2021). These trait values (low SLA, high seed mass and height) are also known to be linked to less intensive agriculture (Fried et al., 2012; Richner et al., 2015). In our models, we observed a similar pattern, with less frequent margin management associated with a decrease along the ruderality axis and an increase along the stress-tolerance axis. All of this suggests that climate change and the evolution towards more extensive agricultural practices will select the same trait values towards more stress-tolerant and less ruderal strategies (MacLaren et al., 2020).

Finally, temporal analyses showed additional phenological changes, suggesting that climate change could increase the occurrence or abundance of late-flowering species, i.e. species that have high temperature requirement to complete their life cycle (Peters et al., 2014). These phenological shifts coincided with a decrease in trait variance, leading to trait convergence within communities. Critically, such changes can reduce the ability of species to escape field margin management, which typically favors species able to flower all-year-round, as expected with a ruderal strategy. As species will not be able to advance their phenology indefinitely, this can ultimately result in species losses in the long-term. However, farmers are likely to adapt the temporality of their practices to climate change, mitigating some of these impacts.

### Agricultural practices have a weaker impact on temporal community dynamics

Temporal variations in agricultural practices over the short-term had a weaker influence on field margin plant communities than climatic variations (Alarcón Víllora et al., 2019; Fried et al., 2019). Field margin management was the most impactful practice, affecting traits related to the ruderal syndrome in a consistent way. This supports the idea that field margin management, as the only practice applied directly in the margin, has a greater impact than herbicides and fertilization applied in the adjacent agricultural fields, which are likely to have only collateral effects.

The lack of herbicide effect on community trait composition could arise because the number of herbicide applications has varied little in recent years or because we have omitted some traits that reflect herbicide tolerance (e.g. leaf cuticle thickness, hairiness). Also, reducing the intensity of agricultural practices may not necessarily influence the trait composition of communities, because agricultural intensification has already greatly reduced functional diversity, and highly diverse landscapes would be required for some species to recolonize field margins.

Fertilization had minimal influence on changes in community trait composition, but reduced species richness (Kleijn & Verbeek, 2000), an effect detected in the MZ and leading to the loss of some annual Mediterranean species (Poinas et al., 2023). Due to functional trade-offs, we saw that nitrophilous plant species were less thermophilous and more acidiphilous, which explains why nitrogen dose was related to affinity for acidic soils in our models. Nitrogen dose remained constant over time, which aligns with the weak change in global nitrophily levels in plant communities, suggesting that eutrophication may no longer be the primary driver of changes in arable vegetation (Alignier, 2018; Duchenne et al., 2021).

### Functional trade-offs and implications for communities response to ongoing global changes

Our findings revealed that resource level (driven by fertilization) and climate vary the position of communities along the stress-tolerance axis (see **Appendix I** for more results on the effect of fertilization), while disturbance level (driven by field margin management) and climate vary the position of communities along the ruderal axis (**Fig. 7**). This supports the view that functional trade-offs are not only evident on a global scale as found by Wright et al. (2004) and Díaz et al. (2016), but also occur within a narrower functional range (such as plants colonizing agricultural field margins). As a result, agricultural intensification and climate change act in opposite ways on the trait composition of field margin plant communities. Climate change favors species that are adapted to high temperatures and drought, but not to intensive agriculture. It tends to expand the functional range for traits related to stress-tolerance within communities (divergence), but reduces the functional range for traits associated to ruderality (convergence). Conversely, agricultural disturbances select species more sensitive to current climatic trends, without any particular trend in trait variance.

**Fig. 7.**
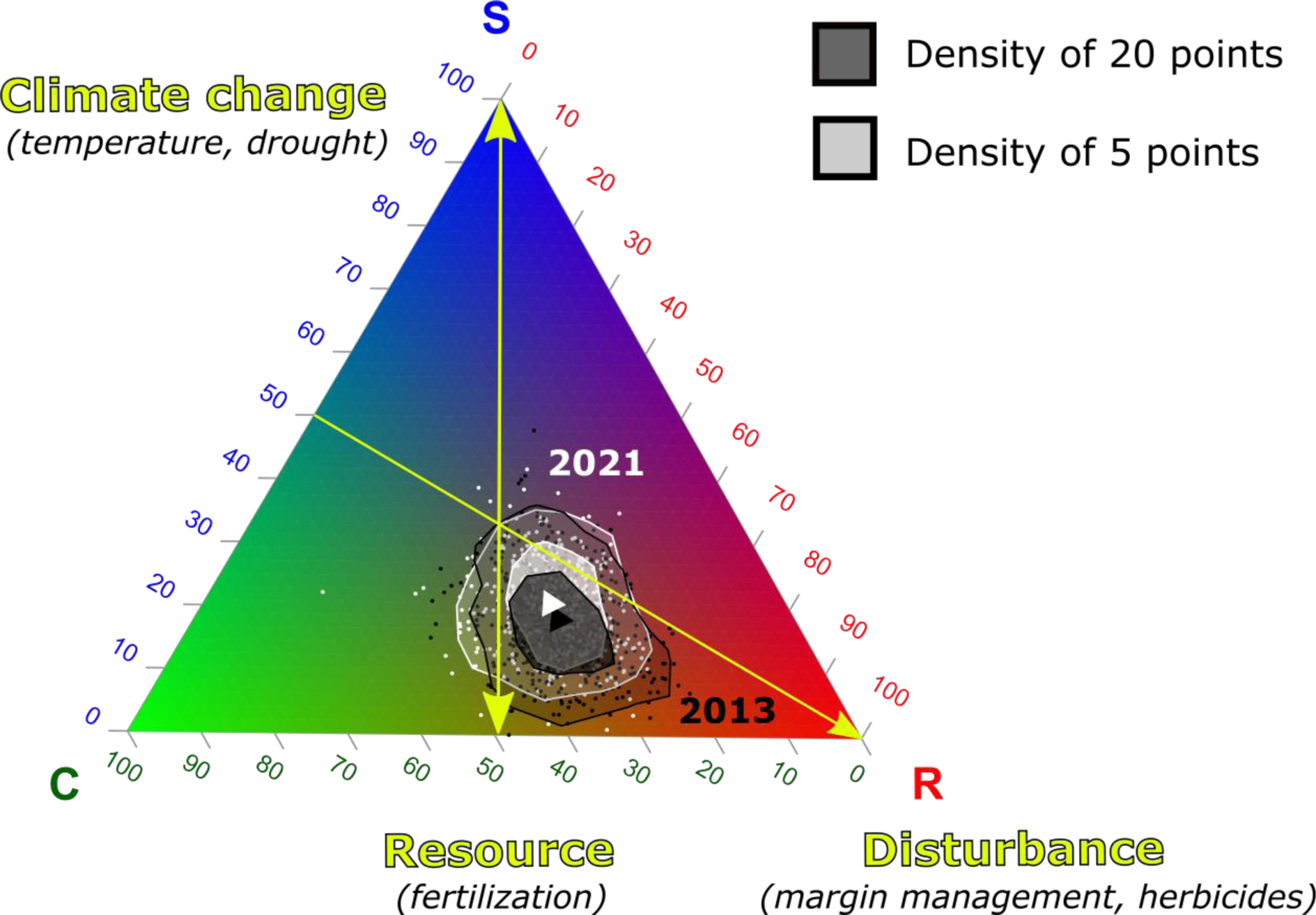
Grime’s CSR triangle depicting the temporal trajectory of community strategies between 2013 and 2021. To enhance clarity, we show only two levels of density curves, with each point representing a specific site. The relative percentages of each strategy are depicted through the use of green, blue and red colors (C = competitor, S = stress-tolerant and R = ruderal). Arrows indicate expected impacts of climate change, disturbance, and resource levels. Temporal models (GAMM) applied to the CWM of each strategy revealed significant decreases in ruderality (−1.14% by decade) and competitiveness (−0.81% by decade) scores of communities, and a significant increase in stress-tolerance scores (+1.80% by decade).

These trade-offs emphasize the need to consider the existing interactions between climate and agricultural practices when predicting future community trajectories (Garnier et al., 2019; Pakeman et al., 2009). Here, we acknowledge the difficulty in quantifying these interactions, particularly given the limited changes observed in practices over time. However, the effects of practices were more perceptible with analyses focused on spatial effects (**Appendix I** and see also Poinas et al., 2023), allowing us to imagine main trends in community trajectories according to several scenarios (**Fig. 7**). Accelerating climate change coupled with an agricultural abandonment and more extensive practices in Europe (Miller et al., 2022; Peeters et al., 2021) will likely result in an increase in xero-thermophilous and conservative species. However, a large part of these species are habitat specialists (e.g. Mediterranean species as found in Munoz et al. (2017); Fried, Chauvel, et al., 2009) and have a high affinity for calcareous soils, which will probably limit their expansion towards the CZ to restricted calcareous areas, such as the Paris Basin. Areas where they are unable to colonize might suffer a decrease in species richness, and this scenario could be worsen if current levels of agricultural intensification are maintained or increased. Mediterranean species expanding in the northern half of France could face severe agricultural intensification that would likely limit their expansion, while at the same time ruderal species would become less frequent mostly because of drought. This highlights the need to consider the conjunction of climate change and intensive agriculture when making future predictions.

## Conclusion

Our study highlights climate as the primary factor affecting field margin plant communities in France, with increasing temperatures and decreasing soil moisture fostering Mediterranean, stress-tolerant and conservative species, while negatively affecting ruderal species. In comparison, agricultural practices had a limited effect on changes in species richness and trait composition, with field margin management having the greatest impact. Our study suggests that the species selected by climate change are poorly adapted to intensive farming, while the pool of species currently able to colonize field margins is restricted to a limited functional range adapted to agricultural practices. The persistence of intensive agricultural practices and accelerating climate change could thus have critical consequences for the conservation of floristic diversity in agroecosystems. However, it is important to consider the potential of adaptation of species, through intraspecific trait variability and phenotypic plasticity (known to be particularly high in ruderal species, Baker, 1974), as it may enhance their resilience to changing conditions. Our findings suggest a reduction in ruderality and an increase in stress-tolerance according to Grime’s strategies. Bopp (2023) highlighted a similar increase of stress-tolerance in weeds in response to climate change, but did not observe a corresponding decrease in ruderality. Further investigations are thus necessary to assess the generalizability of these results across different habitats, including communities with broader or narrower functional niche, such as weeds. Long-term monitoring programs are necessary to address other important research questions, such as the potential time-lag in flora’s response to environmental changes, the non-linearity in temporal trends and the interactive effects between climate and agricultural changes. Finally, the findings presented in this study call for rethinking our current agricultural model, urging us to prioritize the development of agricultural practices that foster the creation of favorable microclimates while minimizing local intensification. Promising approaches, such as agroforestry, hold the potential to align agricultural production with biodiversity conservation goals by providing refuge habitats and microclimate regulation.

## Acknowledgements

The 500-ENI network is developed by the French Ministry of Agriculture under the Ecophyto framework with funding from the French Biodiversity Agency (Office Français de la Biodiversité). We would like to thank everyone that has collected data in the field, the farmers who provided information on their practices, and everyone involved in the coordination of the 500-ENI data network.

Funding: This work was supported by an INRAE-ANSES thesis fellowship, an Ecophyto II+ project: GTP 500 ENI (OFB-21-1642), and the AgriBiodiv ANR-21-CE32-006-01.

## Conflict of interest disclosure

All authors of this preprint declare that they have no financial conflict of interest with the content of this article.

## Appendix A-H. Supplementary data

Supplementary data associated with this article can be found, in the online version, at https://doi.org/10.1101/2023.03.03.530956.

## Notes

https://doi.org/10.57745/QAMAWM

### Competing Interest Statement

The authors have declared no competing interest.

### Summary of Updates

Peer-reviewed and recommended by PCI Ecology

