## Supplementary information for "Functional trade-offs: exploring the temporal response of field margin plant communities to climate change and agricultural practices"

**Appendix A.** Details about the sampling protocol in the 500-ENI network.

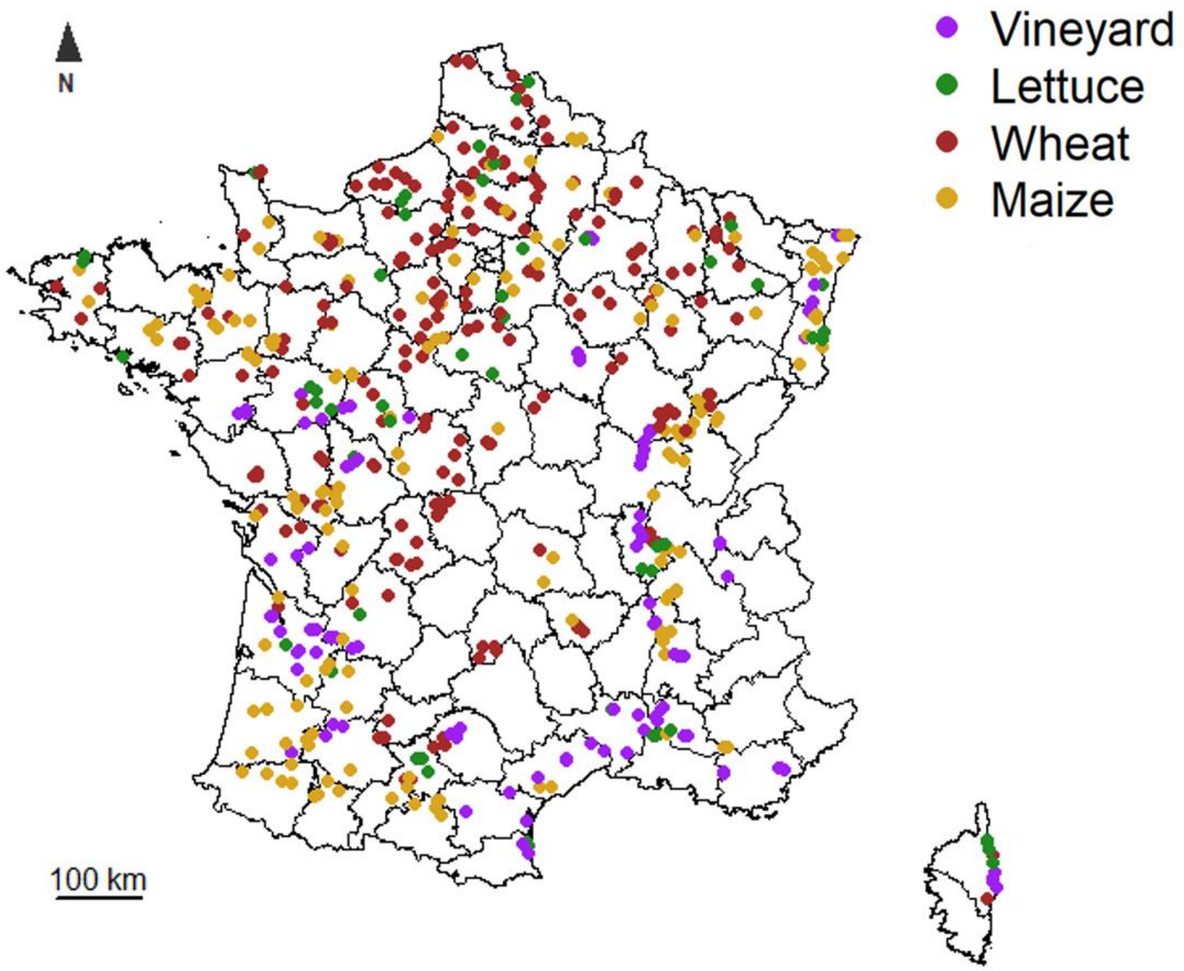


**Fig. A.1.** Distribution map of the 555 field margins monitored at least one year between 2013 and 2021 in continental France. The black lines represent the limits of departments, a French administrative unit dividing continental France into 95 units. Purple: vineyards (n = 93), green: market gardening crops (n = 50), brown: winter wheat in rotation (n = 178), yellow: maize in rotation (n = 141).

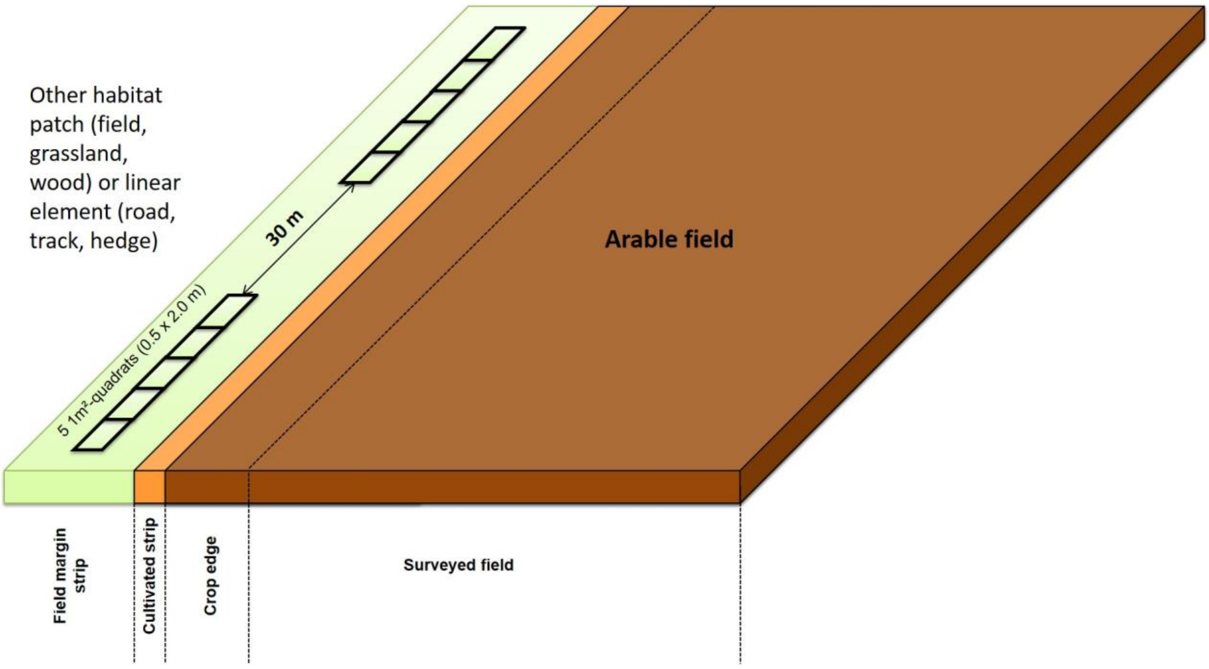


**Fig. A.2.** Details about the sampling protocol within each field margin (Andrade et al., 2021). The *crop edge* is the 1-6 first meters of the crop. The *cultivated strip* or *crop strip* is outside the last row of crops and is mostly composed of bare soil usually colonized by weed species from the field. The *field margin strip* is the uncultivated herbaceous strip between the *cultivated strip* and the *adjacent habitat*.

**Appendix B.** Correlation matrix (with Spearman) of (A) explanatory factors and (B) response variables on the whole dataset. Abbreviations are in **Table 1**.

**(A)**

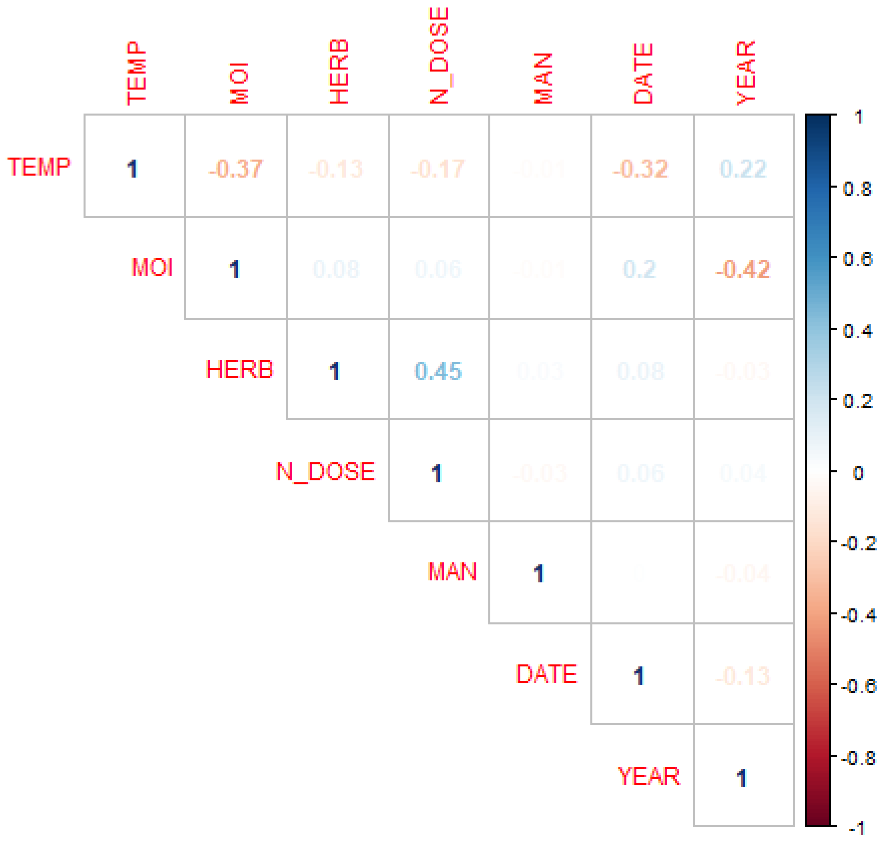

**(B)**

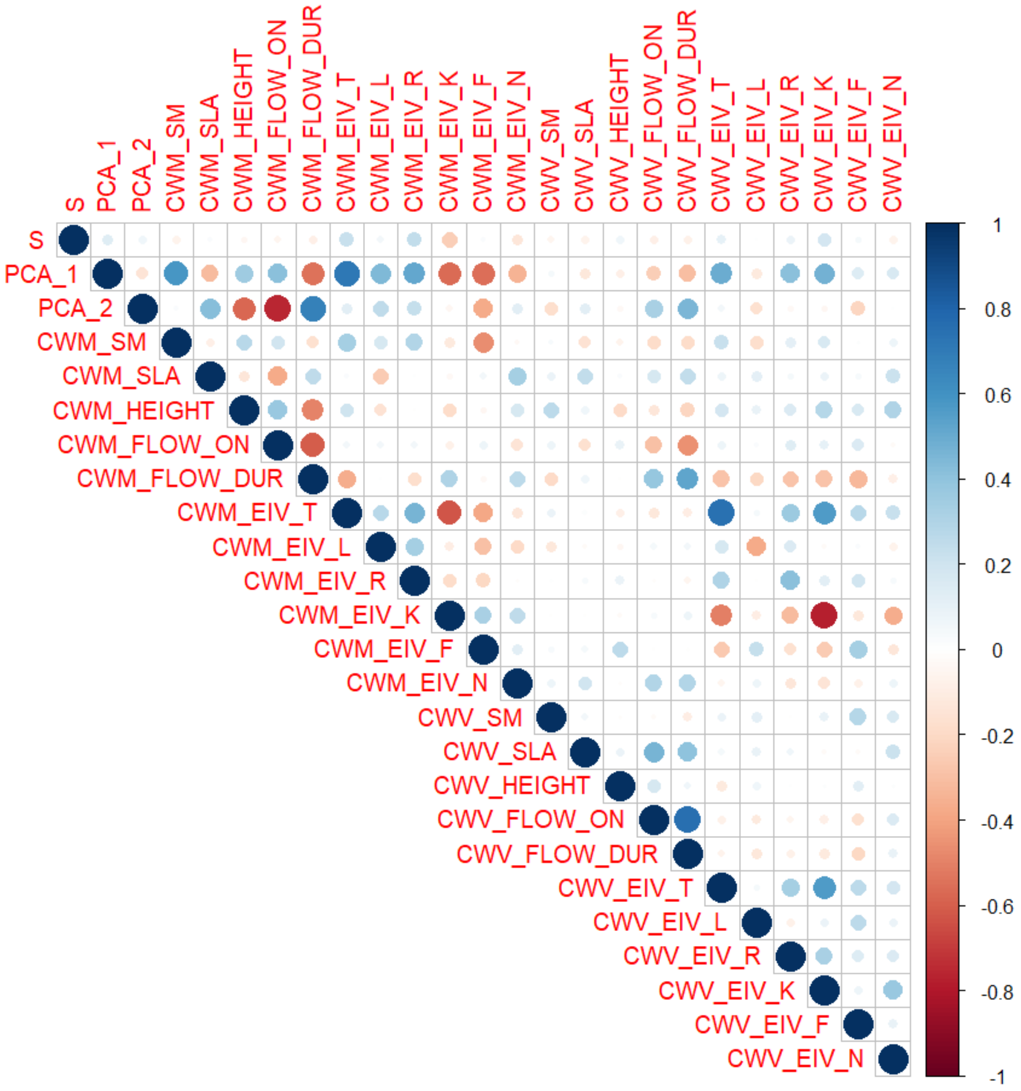
S
PCA_1
PCA_2
CWM_SM
CWM_SLA
CWM_HEIGHT
CWM_FLOW_ON
CWM_FLOW_DUR
CWM_EIV_T
CWM_EIV_L
CWM_EIV_R
CWM_EIV_K
CWM_EIV_F
CWM_EIV_N
CWV_SM
CWV_SLA
CWV_HEIGHT
CWV_FLOW_ON
CWV_FLOW_DUR
CWV_EIV_T
CWV_EIV_L
CWV_EIV_R
CWV_EIV_K
CWV_EIV_F
CWV_EIV_N
1
0.8
0.6
0.4
0.2
0
-0.2
-0.4
-0.6
-0.8
-1

**Appendix C**. Details on the selection of explanatory variables and functional traits, including informations on data extraction and sources. We also include the references used to develop hypotheses of plant response to climatic and agricultural gradients.

For climatic variables, we decided to focus on temperature and soil moisture. Mean annual temperature and precipitation are known to be tightly related to floristic composition and richness in agroecosystems (Fried et al., 2008). However, we focused our attention on soil moisture rather than precipitation, as it is more integrative of the effective water availability to plants and reflects the interaction between temperature and precipitation (Moles et al., 2014). Concerning agricultural practices, we included the number of herbicide treatments, nitrogen dose applied in fertilizers (within adjacent agricultural fields) and the number of margin management events. These variables have consistently been reported to have a significant effect on field margin communities (Aavik & Liira, 2009; Bassa et al., 2011; Fried et al., 2018). Finally, we added the observation date of the floristic survey in temporal analyses, as some studies have reported a seasonal pattern in the succession of herbaceous species (Bopp et al., 2022; Delaney et al., 2015).

For functional traits, we extracted the specific leaf area (SLA) from LEDA (Kleyer et al., 2008), the maximum plant height from Flora Gallica (Tison & de Foucault, 2014), the seed mass from SID (Liu et al., 2019) and the flowering onset, flowering duration, chorology and Ellenberg values from Baseflor (Julve, 2015). SLA relates to the ability to acquire and use resources (Garnier & Navas, 2012), while maximum plant height at maturity reflects the species' competitive ability and depends on resources and disturbances (Fried et al., 2012; Gaba et al., 2014). Seed mass related to the trade-off between competition and dispersal, with small seeds more likely to recolonize after recurrent disturbances in agroecosystems (Fried et al., 2012; Gaba et al., 2014). Flowering phenology (onset and duration) is key in the response of plant species to agricultural disturbances. Early-flowering species with long flowering are more likely to escape management disturbances, as they produce seeds before the first vegetation management and even long after if their flowering period extends after the last intervention (Fried et al., 2012; Fried et al., 2022; Gaba et al., 2017). We added Ellenberg indicator values which estimate the optimal position of a species along particular environmental gradients (Ellenberg, 1974). We retained Ellenberg values for light (L), temperature (T), continentality (K), moisture (F), pH (R) and nutrients (N). Most of these indices are expected to depend on climatic and edaphic factors.

References used to develop ou hypotheses of plant response to agricultural and climatic gradients (see **Table 1** for abbreviations and colors, **R** = resource, **Di** = disturbance, **Dr** = drought, **T** = increasing temperature, **(+)** = positive relationship, **(-)** = negative relationship, **(+/-)** = relationship with contradictory observations in the literature):

- DATE: **R(-)** / **Di(-)** (Bopp et al., 2022)
- SLA: **R(+)** / **Di(+)** (Borgy et al., 2017; Garnier & Navas, 2012; Kazakou et al., 2016; Wheeler et al., 2023)

  **Dr(-)** (Garnier et al., 2019; Kühn et al., 2021; Moreau et al., 2022)

  **T(+/-)** (Alarcón Víllora et al., 2019; Kühn et al., 2021)
- HEIGHT: **R(+)** / **Di(-)** (Gaba et al., 2014; Wheeler et al., 2023)

  **Dr(+/-)** (Garnier et al., 2019; Kühn et al., 2021; Moreau et al., 2022)

  **T(+/-)** (Alarcón Víllora et al., 2019; Kühn et al., 2021)
- SM: **R(+)** / **Di(-)** (Gaba et al., 2014; MacLaren et al., 2020; Pakeman et al., 2009)

  **Dr(+)** (Alarcón Víllora et al., 2019; Cochrane et al., 2015; Garnier et al., 2019)

  **T(+)** (Thuiller et al., 2005)
- FLOW_ON: **Di(-)** (Gaba et al., 2017; MacLaren et al., 2020)

  **Dr(+)** / **T(+)** (Nordt et al., 2021; Peters et al., 2014)
- FLOW_DUR: **Di(+)** (Fried et al., 2022; MacLaren et al., 2020)

Dr(-) / T(-) (by opposition to flowering onset)

- EIV_N: R(+) (Fried et al., 2009)
- EIV_F: Dr(-) (Stanik et al., 2021)
- EIV_T: Dr(+) (Stanik et al., 2021)

  T(+) (Fried et al., 2020; Martin et al., 2019)
- S: R(+/-) (Grime, 1988; Tilman, 1987)

  Di(+/-) intermediate disturbance hypothesis (Connell, 1978 *in* Bassa et al., 2012)

  Dr(+/-) / T(+/-) (Duchenne et al., 2021; Thuiller et al., 2005)
- CWM_C : R(+) / Di(-) / Dr(-) (Grime, 1977, 1988)
- CWM_S: R(-) / Di(-) / Dr(+) (Grime, 1977, 1988)
- CWM_R: Di(+) (Grime, 1977, 1988)

**Appendix D.** Null models to correct for CWV dependence on species richness.

Randomization was repeated 999 times to produce a random distribution of CWV values. From these simulated values, we computed an effect size (ES) with the following formula:

$$ES = \frac{number\ (null\ < obs)}{999} - 0.5$$

Where *number (null < obs)* is the number of simulated CWV out of 999 that are smaller than the observed CWV. Positive ES values denote a convergence in the trait values within the community, while negative ES values indicate a divergence. We chose this form of standardization over Standardized Effect Sizes, because some of our null distributions were not normal and skewed (Bernard-Verdier et al., 2012; Perronne et al., 2017).

**Appendix E.** Details about distribution laws and removed values in response variables of temporal models (GAMMs). Some values of CWM of Ellenberg-K were removed as their frequency far exceeded that of all others, thus yielding to a non-Gaussian distribution.

| | | Removed values | Distribution law of the temporal model (GAMM) |
|---|---|---|---|
| | **Species richness** | | Negative binomial |
| | **PCA Axis 1** | | Gaussian |
| | **PCA Axis 2** | | Gaussian |
| | **Seed mass** | | Gamma (link = log) |
| | **Specific leaf area** | | Gaussian |
| | **Maximum plant height** | All values superior to 4m (two creepers species) These values were directly removed within traits and not CWM. | Gaussian |
| **CWM** | **Flowering onset** | | Gaussian |
| | **Flowering duration** | | Gaussian |
| | **Ellenberg-T** | | Gamma (link = log) |
| | **Ellenberg-L** | | Gaussian |
| | **Ellenberg-R** | | Gaussian |
| | **Ellenberg-K** | All values equal to 5 (intermediate continentality) | Gaussian |
| | **Ellenberg-F** | | Gaussian |
| | **Ellenberg-N** | | Gaussian |
| | **All CWVs** | | Gaussian |

**Appendix F.** Estimates of the temporal models (GAMM) of each predictor with the year as the only explanatory factor. Note that estimates are computed for a decade and are only comparable for the same predictor. In row, the different subsets of data (see **Fig. 1**) and in column, the different factors. Non-significant coefficients are noted as 0.

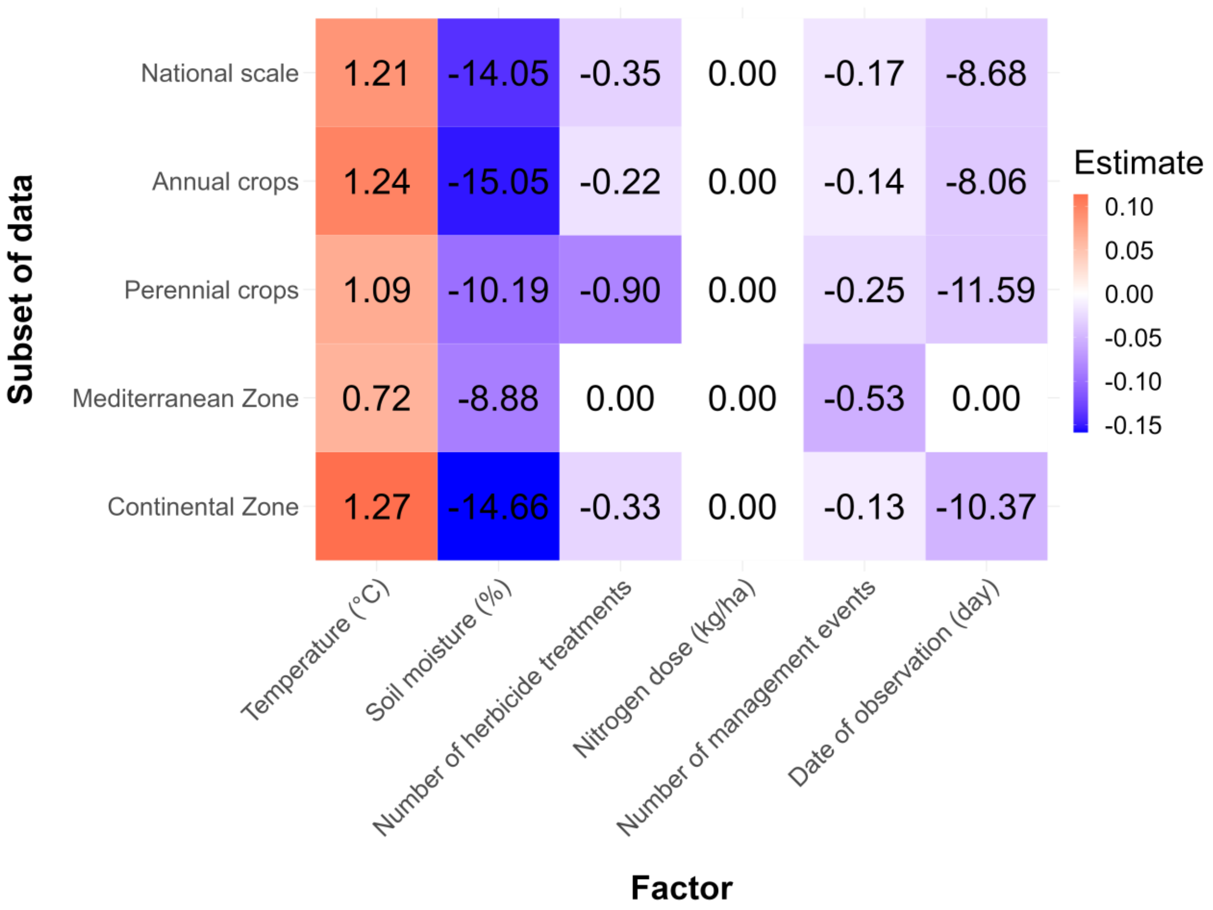

**Appendix G.** Details about PCA on CWM of functional traits and PCA on functional traits of species. Abbreviations are in **Table 1**.

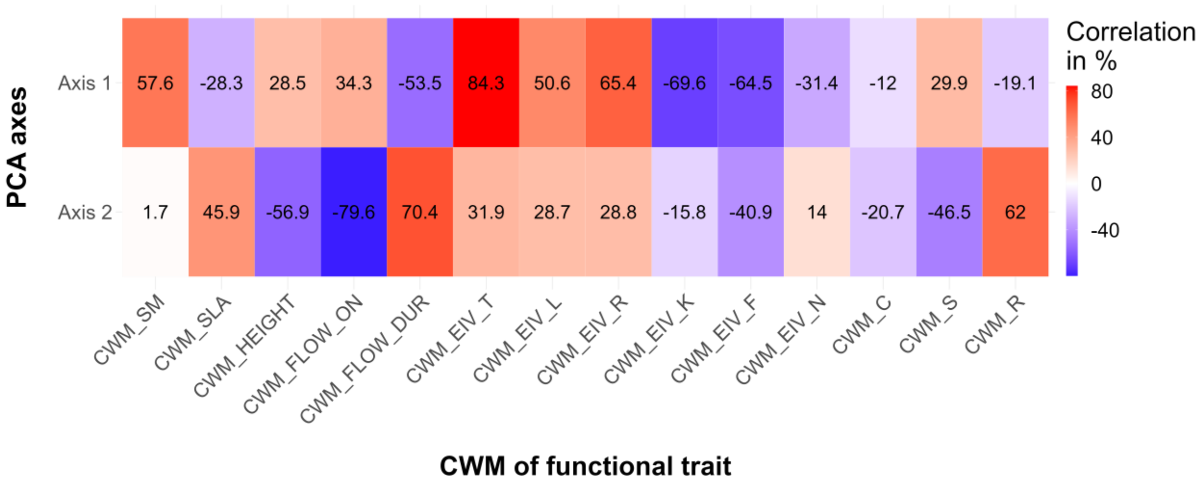


**Fig. G.1.** Correlation of each functional trait and strategy to PCA axes (**Fig. 4**), for PCA on CWM of functional traits computed by observation.

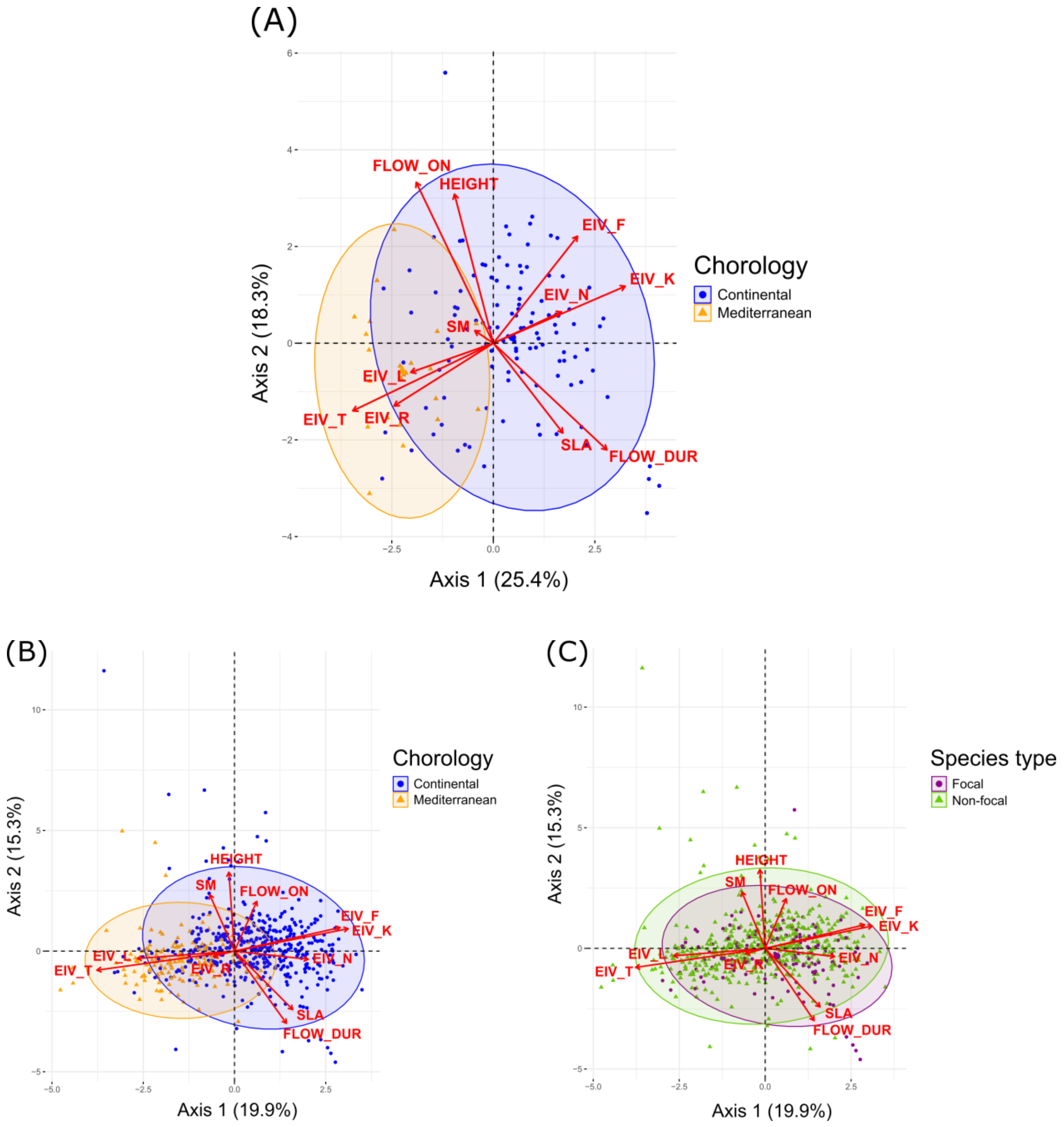


**Fig. G.2.** Normed PCA on functional traits of the (A) 140 focal species and (B-C) 601 species for which we have all the traits. The chorology of species was extracted from Baseflor (Julve, 2015) and is depicted by colored ellipses in (A) and (B). In (C), the set of focal species is compared to the set of non-focal species. The difference between the two sets of species was assessed with Wilcoxon tests and was significant for the axis 1 ($p = 0.001$) and 2 ($p = 0.003$). However, the results of temporal models, on the entire species dataset did not differ from those obtained with the focal species (not shown here).

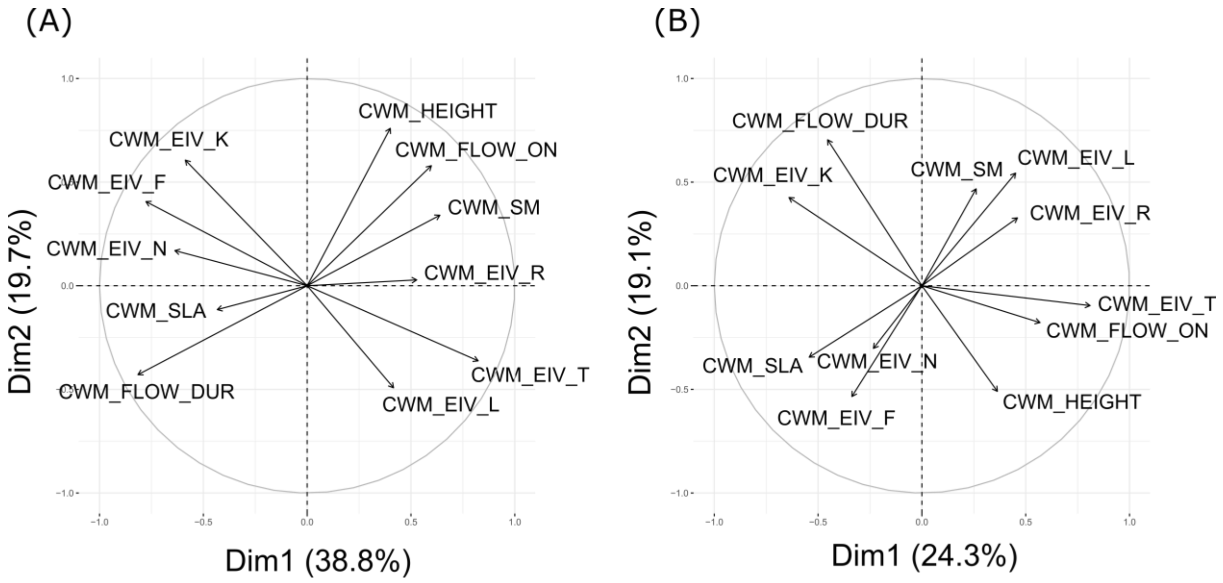


**Fig. G.3.** Normed PCA on CWM of functional traits computed by observation (first two axes), as in the **Fig. 4**. (A) For annual species. (B) For perennial species.

**Appendix H.** Results of temporal models (GAMM) implemented on each response variable (in rows) as illustrated in **Fig. 5,** for different subset of data (**Fig. 1**). Explanatory factors are in columns. The legend indicates the corresponding order of cells with the order of data subsets. Trait names refer to the CWM (results on CWV not shown here). Colored cells are the significant estimates (p-value threshold of 0.01). Positive estimates are in red, negative estimates in blue, and the strength of the relationship (based on the standardized estimates) is reflected by the lightness of the color (weaker when lighter).

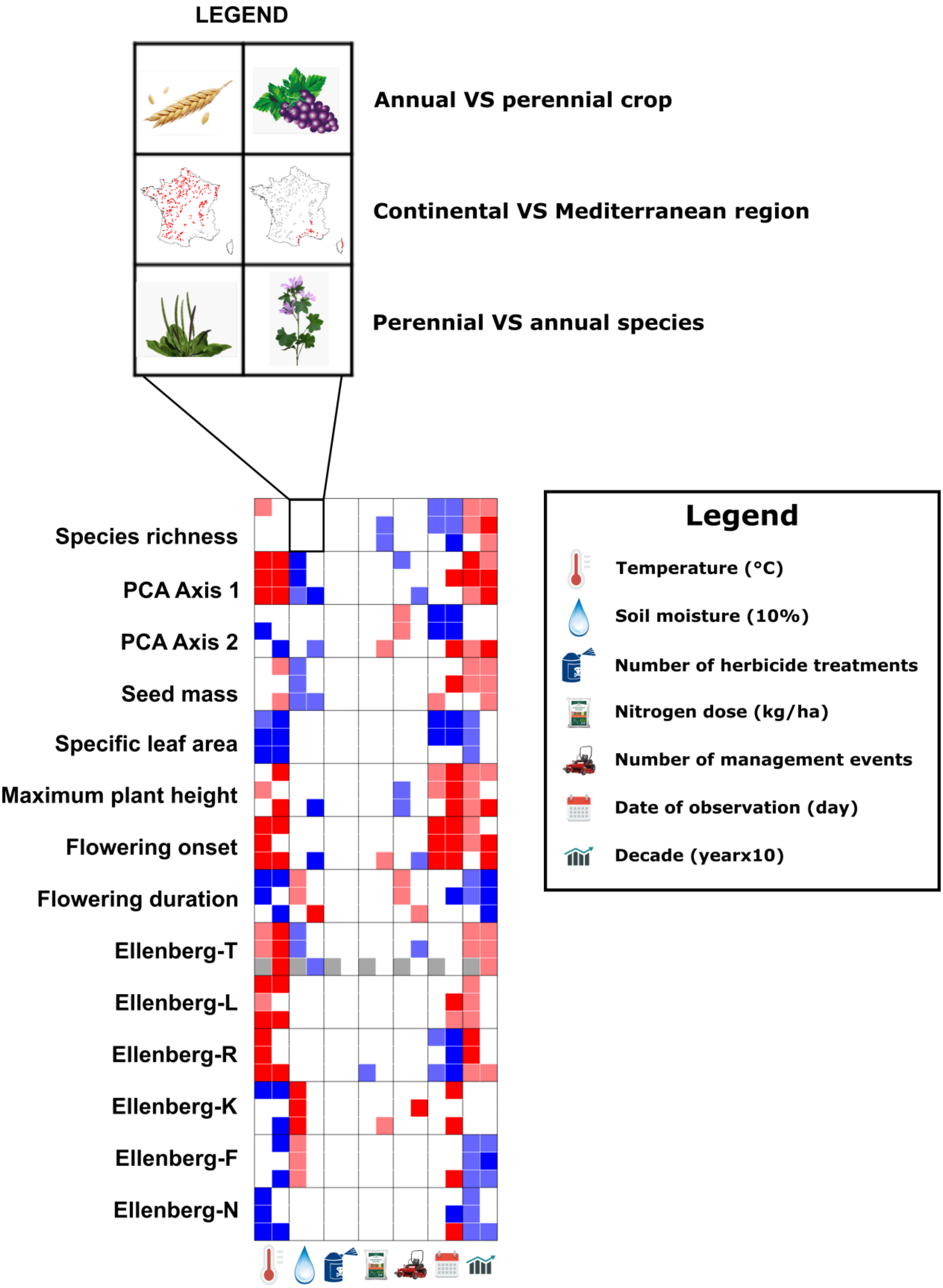
LEGEND
Annual VS perennial crop
Continental VS Mediterranean region
Perennial VS annual species
Species richness
PCA Axis 1
PCA Axis 2
Seed mass
Specific leaf area
Maximum plant height
Flowering onset
Flowering duration
Ellenberg-T
Ellenberg-L
Ellenberg-R
Ellenberg-K
Ellenberg-F
Ellenberg-N
Legend
Temperature (°C)
Soil moisture (10%)
Number of herbicide treatments
Nitrogen dose (kg/ha)
Number of management events
Date of observation (day)
Decade (yearx10)

**Appendix I.** Methodology and results for spatial models implemented on the same dataset as temporal models.

The purpose of these models was to ignore temporal trends and variations to examine the spatial response of plant communities (in terms of species richness, trait composition and ecological strategies) to climatic and agricultural factors. With these spatial models, we aimed to reveal agricultural effects that may not be detected through temporal analyses. This approach allows us to determine whether the absence of these temporal effects is real or is the consequence of a lack of clear temporal trends in these agricultural factors.

*Data analysis*

To analyze the effects of spatial variations in climate and agricultural practices while ignoring temporal patterns, explanatory factors and species abundances were averaged across years within sites having at least five years of data, leaving a total of 349 sites. Spatial simultaneous autoregressive models (SAR; package SPDEP, function *errorsarlm*; Cressie, 2015) were implemented to model linear relationships that take into account spatial autocorrelation in the data, i.e. the tendency of nearby points to have more similar values than expected by chance. We examined the relationship between each response variable (species richness, trait composition, CWV and strategies) and the explanatory factors (temperature, soil moisture, nitrogen dose, herbicides and margin management). The Nagelkerke pseudo-$R^2$ (which can be interpreted similarly to a conventional $R^2$) was used to assess the model's explanatory power, and we controlled for the observer bias by adding the number of successive observers in a site as a fixed effect. To compute the CWVs by site, the species pool was defined by the biogeographic region, allowing us to examine spatial variations. To determine the biogeographic regions of each site, we used the VégétalLocal map (Office français de la biodiversité, 2021).

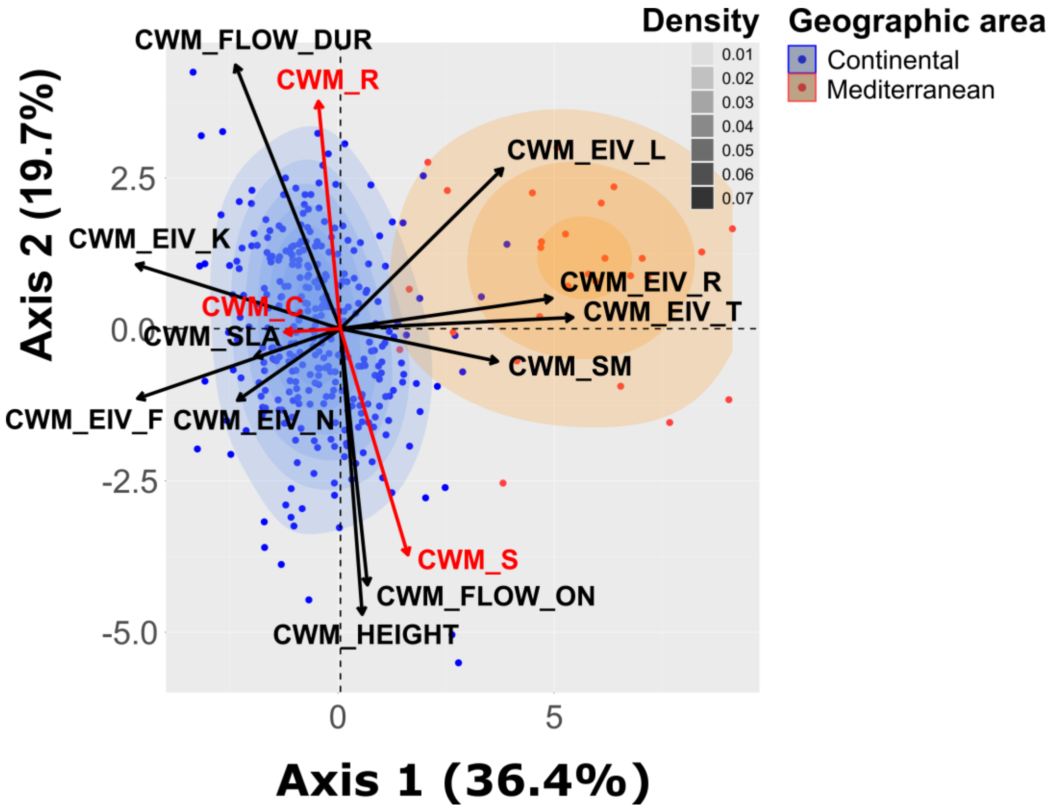


**Fig. I.1.** Normed PCA on CWM of functional traits computed by site. The color of the dots indicates the region to which they belong and the density curve highlights the concentration of data points in a given area. The CWM of strategies (in red) were added as supplementary variables. Abbreviations for CWM: CWM_SLA, specific leaf area; CWM_HEIGHT, maximum plant height; CWM_SM, seed mass; CWM_FLOW_ON, flowering onset; CWM_FLOW_DUR, flowering duration; CWM_EIV_L/T/K/F/R/N, requirement for light/temperature/continentality/moisture/pH/nitrogen; CWM_C, competitive strategy; CWM_S, stress-tolerant strategy; CWM_R, ruderal strategy.

*Results of spatial analyses of plant communities*

Spatial models revealed that climate had a predominant impact on community trait composition and particularly on Ellenberg values, opposing in a consistent way Mediterranean communities to nitrophilous continental ones along the stress-tolerance axis (**Fig. I.1**). Temperature increased the

CWM and CWV of seed mass (divergence) and decreased the CWM and CWV of SLA (convergence). High temperatures lead to more divergence in all environmental requirements (compared to a community of equal richness), except for the requirement for light (Ellenberg-L). Conversely, soil moisture brought convergence in the requirement for temperature and continentality (Ellenberg-T and K). Field margin management favored ruderal communities with higher SLA and/or lower seed mass (PCA Axis 1), shorter stature (-5.2 cm by management event) and longer flowering duration (+3.7 days by management event; PCA Axis 2). Herbicide applications had no significant effect, while the average annual nitrogen dose in fertilizers slightly decreased the species richness ($-3.2 \times 10^{-2}$ species by kg/ha) and pH requirement, and increased the nitrogen requirement, SLA and divergence in flowering duration (**Fig. I.2**). The number of observers surveying a site over the 9-year period was positively correlated with species richness (average increase of 1.7 species by observer) and with the CWM and CWV of flowering duration (divergence).

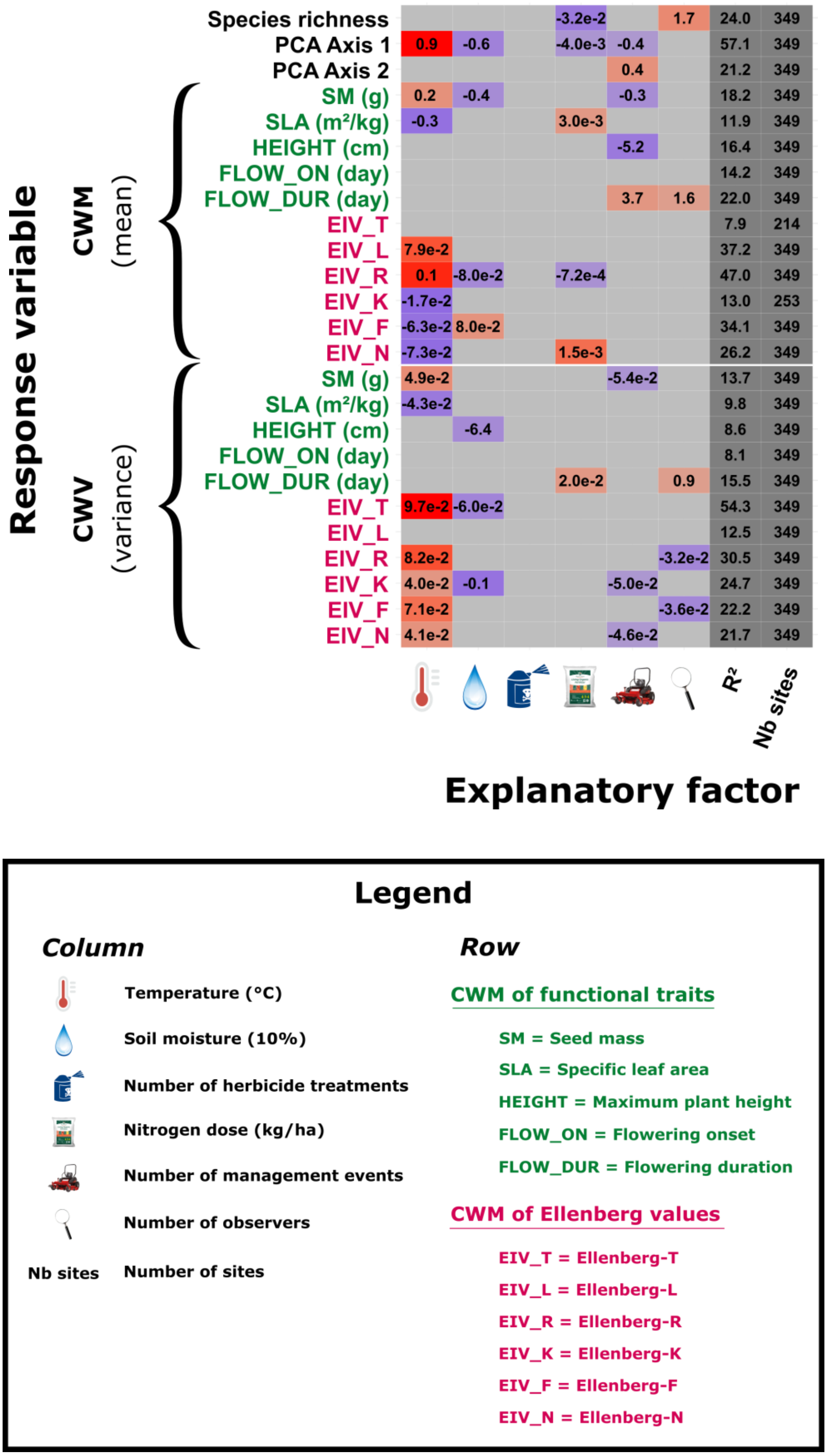


**Fig. I.2.** Results of spatial models (SAR) on the whole dataset, with response variables in rows and explanatory factors in columns. The adjusted R², expressed as a percentage of variation, and the number of observations are reported. Significance is indicated by colored cells, with a p-value threshold of 0.01. Positive estimates are in red, negative estimates in blue, and the strength of the relationship (based on the standardized estimates) is reflected by the lightness of the color (weaker when lighter). It is important to note that the strength of the relationship can only be compared among explanatory factors for a same response variable. Reported values are the raw estimates and can be interpreted in the units of response and explanatory variables (e.g. an increase of 1°C in temperature leads to an increase in 0.2 g in the CWM of seed mass).

Office français de la biodiversité, 2021. Carte des régions d'origine - Référentiel technique de la marque Végétal local - Version 2021.
